# How Similar Are Two Brains? A Comprehensive Benchmark of Brain Network Similarity Measures

**DOI:** 10.64898/2026.09.16.752009

**Authors:** Adrian Dendorfer, Andrea I. Luppi, Francesco Poli, Alexa Mousley, Duncan E. Astle, Kayson Fakhar

**Affiliations:** MRC Cognition and Brain Sciences Unit, University of Cambridge, Cambridge, UK; Technical University of Munich, Germany; Department of Psychiatry, University of Oxford, Oxford, UK; Division of Information Engineering and St John’s College, University of Cambridge, Cambridge, UK; Montréal Neurological Institute, McGill University, Montréal, QC, Canada; Department of Psychiatry, University of Cambridge, UK; Institute of Computational Neuroscience, University Medical Center Hamburg-Eppendorf, Germany

## Abstract

A central goal of neuroscience is to establish how similar (or different) brain networks are across individuals, development, psychiatric conditions, and even between biological and artificial brains. Whenever one such comparison is made, it relies on the assumption that we have a measure that produces an accurate and plausible similarity score. Yet, the field lacks one such measure, and the wide variety of existing measures often produces conflicting results. To address this problem, we perform a systematic comparison of 16 established similarity measures. First, we show that different measures consistently disagree on which brain networks are most similar. Second, we rank all measures according to multiple criteria such as biological plausibility, computational efficiency, and sensitivity. We show that DeltaCon is the most accurate measure in terms of parameter recovery, and that it ranks among the fastest and most noise-tolerant. For this reason, we conclude that DeltaCon is the best general-purpose similarity measure. However, no measure ranks best on more than two of the five criteria we assessed, and different measures should be preferred depending on the exact research question. Measure selection can now be made on evidence and should be a crucial decision in any study that compares brain networks.

## 1 INTRODUCTION

The connectome, i.e. the brain’s wiring network, defined by its regions (nodes) and the white matter tracts that link them (edges), is highly organized^1^. Its topology exhibits complex features, some shared with other complex networks^2^ such as with power grids^3^, the internet^4^, and metabolic networks^5^. This shared organization means the brain can be analysed with the tools of network science, and studying its topological features yields insights into the connectome’s behavior, including its response to perturbation^6^. Research on connectomes has also revealed that many neurological and psychiatric conditions can be characterized by connectome abnormalities and altered topological features^7,8^. One approach to studying these organizational principles is through computational modelling of network formation.

Generative network models (GNMs) represent one such framework^9,10^. They ask which simple rules suffice to reproduce brain-like wiring: rather than simulating biological growth per se, they compress connectome organization into a handful of interpretable parameters. Concretely, a GNM grows a synthetic network edge by edge, adding connections according to rules that capture the trade-offs a developing brain faces. Biological connections consume space and metabolic energy; this wiring cost must be offset by some wiring benefit, best operationalized as homophily - the principle, observed across many real-world networks, that nodes with similar connectivity profiles (those sharing many neighbours) are more likely to become connected. In the model, wiring cost (controlled by parameter *η*) penalizes long-range connectivity based on Euclidean distance, reducing wiring probability as distance increases, while *γ* scales how strongly shared neighbours are rewarded^9^. This formulation has demonstrated a better fit to empirical brain networks than alternatives driven by spatial distance alone or by node degree^11,12^. A GNM is fitted to a given brain by sweeping these two parameters, generating a synthetic network at every combination, and keeping the combination whose network comes closest to the empirical one - which requires a way of measuring how close two networks are. Applied to empirical data, these models have proven powerful. For instance, wiring parameters track neurodevelopmental trajectories (e.g. age, gray matter morphology, and cognitive scores) at the individual level^12^ and relate to individual differences in cognitive performance and indicators of mental health disorders^13^. All such findings, however, depend on the measure that was used to select the parameters in the first place.

Comparing brain networks is fundamental to network neuroscience well beyond the fitting of generative models. Pairwise comparisons of empirical connectomes have, for example, revealed that network similarity recapitulates mammalian taxonomic order and correlates with phylogenetic distance^14^, and uncovered social relationships within human communities on the basis of functional connectome similarity^15^. Yet whether between two empirical networks or between an empirical and a synthetic one, comparison is not a uniquely defined operation: the answer depends on what notion of similarity one chooses. Two randomly wired graphs, for instance, may appear nearly identical when both are treated as instances of a random network model, yet share almost no edges in common.

The challenge of selecting the right comparison operation is not unique to network *neuro*science. Across the broader field, methods for comparing networks range from edge-wise approaches (such as correlating adjacency matrices) to global structural summaries (like portrait divergence, which captures differences in shortest-path distributions via Jensen-Shannon divergence^16^, or DeltaCon which quantifies differences in node affinities through belief propagation^17^). Only some require the two networks to share a node set, which here they do. Network comparison methods have been benchmarked systematically in general settings^18–21^, and those benchmarks show that the choice of measure can substantially change which networks are judged similar. For brain networks, however, no equivalent evaluation exists to our knowledge: prior work has compiled lists of measures used for brain network comparison without empirically comparing them, explicitly raising the need for validation using simulated data^22^.

Within network neuroscience specifically, the field has converged on ”KS-based energy”, a distance measure defined as the maximum Kolmogorov-Smirnov (KS) distance among the four graph-theoretic properties degree, clustering coefficient, betweenness centrality, and edge length^11^. Its appeal is that it is transparent about what it compares: the parameter sweep described above minimizes a quantity built from properties neuroscientists already interpret. Recently, however, this measure was found to have limitations: models fitted by minimizing it systematically fail to reproduce the anatomical placement of long-range connections^23^, potentially contributing to the selection of models that are topologically brain-like - reproducing *who* connects to whom - but topographically very different from the brain, placing those connections in the wrong anatomical locations^24^.

Together, these considerations highlight the pressing need to evaluate the legitimacy of the evaluator itself. To this end, we benchmarked 16 distance measures drawn from different areas of network science to establish principled guidelines for brain network comparison. We used GNMs as a controlled experimental platform: generating 25,000 networks by sweeping through the two parameters *η* and *γ* provides systematic coverage of network space and enables ground-truth parameter recovery experiments unavailable with empirical data alone. The GNM parameter space can be framed as a morphospace^25,26^, i.e. a geometric hyperspace whose axes are model parameters (here *η* and *γ*) and whose regions correspond to networks with distinct morphological characteristics. Within this morphospace, we computed the distance of every generated network to an empirical human consensus connectome, i.e. a group-level network retaining the connections found consistently across subjects, and identified the parameter region most closely resembling empirical connectomes. We structured our analysis around five complementary axes: First, we examined **agreement**: did different measures agree on which networks were similar, or did they identify entirely different networks as maximally brain-like? Second, we investigated **biological plausibility**: did measures identify parameter regions producing networks that satisfied basic biological constraints, such as the penalisation of metabolically costly long-range wiring together with realistic degree and connection-length distributions? Third, we assessed **computational efficiency**: how did measures differ in speed, and what performance trade-offs accompanied faster algorithms? Fourth, we evaluated **sensitivity and robustness**: how did a measure respond to topological perturbations of different magnitudes, and is its result robust to noise in the networks it compared and in the reference it was fitted to? Fifth, we assessed **accuracy**: did measures correctly recover known generative parameters when the true answer was available, and did they rank real connectomes ahead of synthetic ones? Three of the experiments in this benchmark are checked against a known answer: the parameter recovery experiment knows the parameters each test network was generated from, the real-versus-synthetic test knows which networks came from a brain, and the rewiring experiment knows exactly how many edges were moved away from the reference network, so a measure’s response can be held against the magnitude of perturbation that the network experienced.

No measure leads on more than two of the five axes, so the choice between them is a trade-off rather than a ranking. Figure 1D condenses the axes into one spoke each of a per-measure profile, so those trade-offs can be read at a glance. Which measure a study uses therefore determines which networks it calls brain-like.

**Figure 1:**
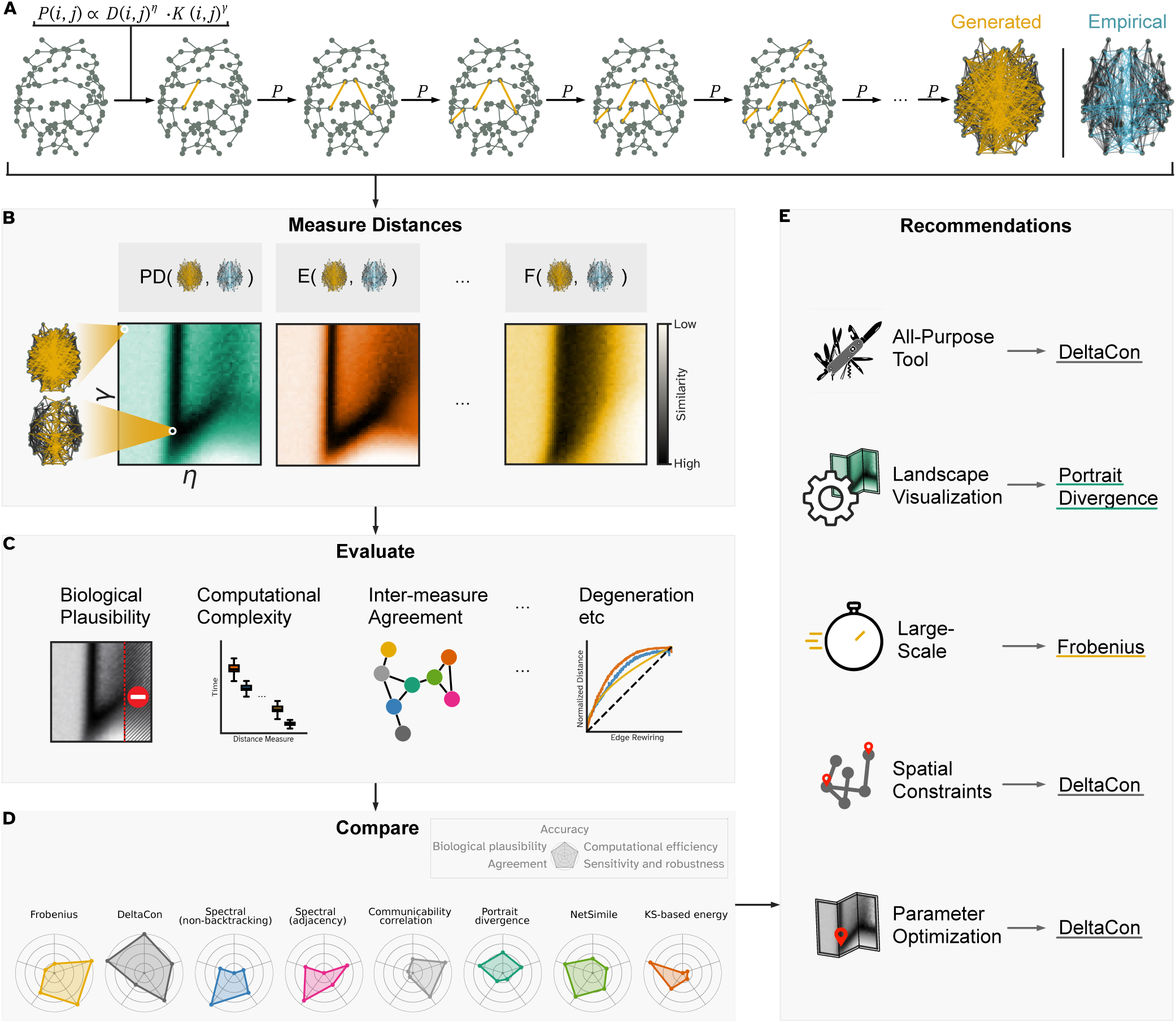
Study overview. **A** A generative network model (GNM) adds connections one at a time, each drawn from a wiring probability *P* (*i, j*) that trades the wiring cost between two nodes (*D*, weighted by *η*) against their homophily (*K*, weighted by *γ*), until the generated network (orange) reaches the density of the empirical connectome (blue; 495 edges, 10% density). A 50 × 50 sweep of *η* and *γ* with 10 replicates per combination gives a morphospace of 25,000 networks. **B** Each of 16 distance measures scores every generated network against an empirical human consensus connectome, giving one landscape per measure. Three are shown (portrait divergence, KS-based energy, Frobenius), with *η* on the horizontal and *γ* on the vertical axis and darker colours closer to the empirical connectome. **C** Four of the evaluations applied to the measures: biological plausibility, computational complexity, inter-measure agreement, and the response to progressive edge rewiring. The full set is reported in the Results. **D** One radar plot per selected measure, with one spoke per evaluation axis. Each spoke carries the measure’s rank among the eight on that axis, so a larger radius is always the more desirable value. Rings mark ranks 2, 4, 6, and 8. Accuracy is scored differently: a measure whose recovery error reaches the chance level of the window sits at 0, and the measures better than chance are spread linearly between the centre and the best error. The read-out behind each axis is given in Section 4.13. **E** Recommended measure for five use cases: an all-purpose default, landscape visualization, large-scale application, spatial constraints, and parameter optimization. The reasoning is given in the Discussion.

## 2 RESULTS

### The morphospace

For an overview of this work see Figure 1. Networks are generated by sweeping the two parameters of a generative network model (GNM), *η* and *γ*, over a 50 × 50 grid with 10 replicates per combination. This yields the 25,000 networks used throughout; the networks and their underlying structure are the *morphospace*. A visualization of how the GNMs are generated is in Figure 1A, more information on GNMs is in 4.2.

### Distance Measures

We evaluated 16 distance measures spanning multiple methodological categories (see Section S1 for full definitions; for visualizations of the mechanisms of the most important measures see Figure 2 and Section S1). *Matrix-based measures* compare adjacency matrices directly: Frobenius distance (Euclidean norm of the element-wise difference), Hamming distance (count of differing edge positions^27^), Jaccard distance (one minus the ratio of shared to total edges^28^), and F1 distance (one minus the harmonic mean of edge-overlap precision and recall^29^). *Information-theoretic measures* include Network Mutual Information (NMI; one minus the normalized Shannon mutual information between edge-level distributions of two graphs^30^) and its Degree-Corrected variant (DC-NMI), which computes per-node mutual information to account for degree heterogeneity^30^. *Spectral measures* compare eigenvalue spectra: spectral distance (adjacency)^31^ and spectral distance (normalized Laplacian)^31^ use the Euclidean distance between eigenspectra of the respective matrices, while spectral distance (non-backtracking)^32^ uses the Wasserstein distance between spectra of non-backtracking matrices, which encode paths that never immediately reverse direction. *Graph kernel and feature-based measures* include portrait divergence (Jensen-Shannon divergence between network portraits, which encode the full distribution of shortest-path lengths^16^), NetSimile (Canberra distance - a weighted *L*_1_ distance - between per-node feature vectors aggregating degree, clustering, and egonet statistics^33^), and DeltaCon (Matusita distance - a root-Euclidean distance - between node-influence matrices derived from fast belief propagation^17^, capturing both local and global differences). *Communicability-based measures* operate on matrix exponentials *e^A^*: communicability correlation (Pearson correlation between vectorized *e^A^*) and communicability JSD (Jensen-Shannon divergence between normalized communicability distributions^34^); resistance distance computes the *L*_2_ norm of the difference between effective-resistance matrices, derived from the Moore-Penrose pseudoinverse of the graph Laplacian^35^. Finally, the established *KS-based energy distance*^11^ serves as the baseline, combining Kolmogorov-Smirnov statistics across four network property distributions: degree, clustering coefficient, betweenness centrality, and edge length. All measures were computed on binarized undirected graphs; detailed mathematical definitions are in Section S1. Five of the 16 measures return a similarity rather than a distance: Jaccard, F1, communicability correlation, NMI, and DC-NMI. Every one of them is oriented into a distance immediately after its calculation by negation. Thus, all quantities reported in this work are distances, for which a smaller value always means more similar and a positive correlation between two measures always means agreement.

**Figure 2:**
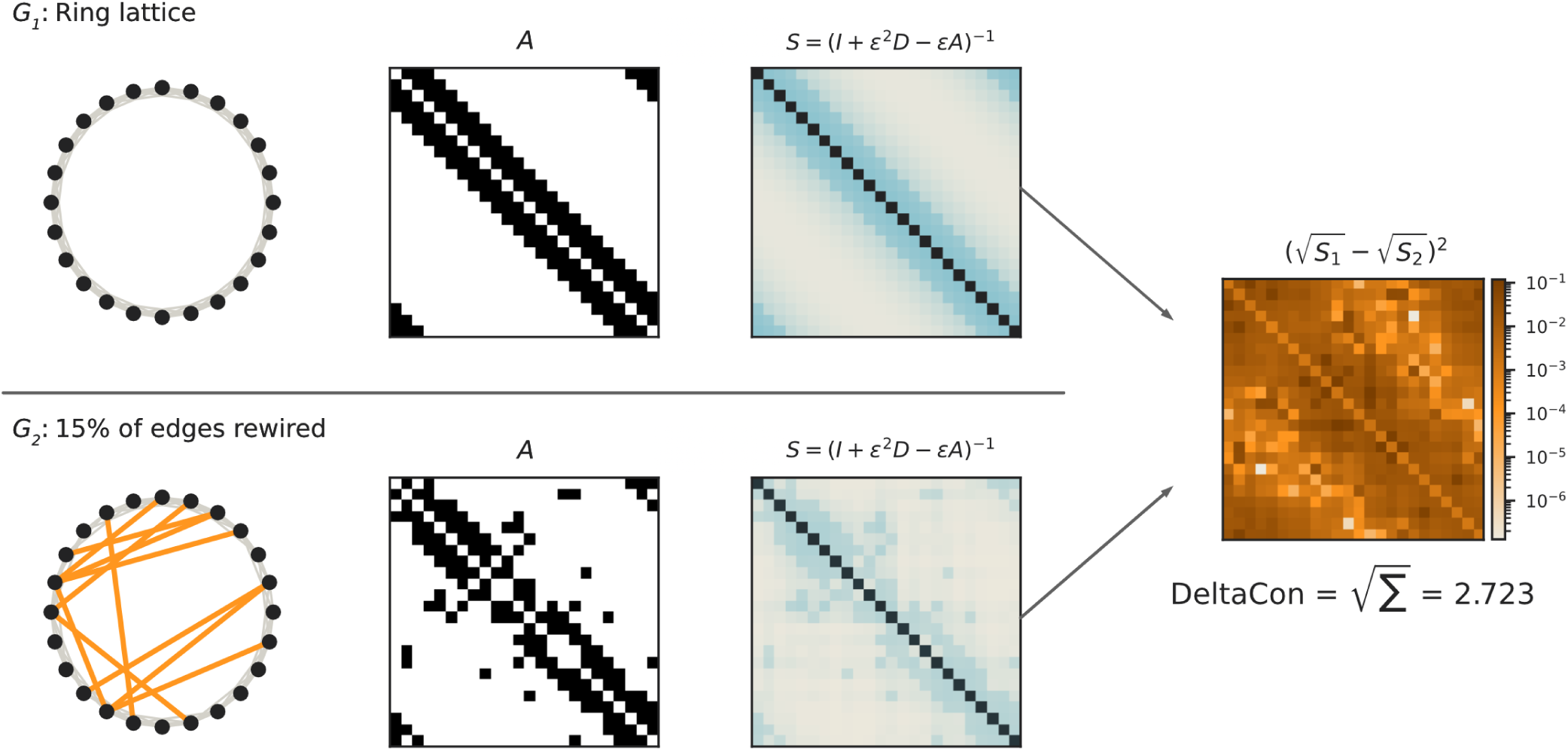
Mechanics of DeltaCon, illustrated on one pair of graphs. *G*_1_ is a ring lattice of 24 nodes in which every node is joined to its three nearest neighbours on either side; *G*_2_ is that same lattice with 15% of its edges rewired to random endpoints, drawn in orange. Both graphs have the same number of nodes and edges, so they differ only in where the connections sit. Per row: the graph, its adjacency matrix *A*, and the fast belief propagation matrix *S* = [*I* + *ɛ*^2^*D − ɛA*]^−1^, whose entries give the influence between each pair of nodes. Right: the element-wise term 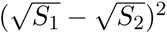 of the Matusita distance on a logarithmic colour scale. The value printed below is the square root of the sum over all entries. The equivalent figures for the other seven selected measures are given in Section S1.

### Distance landscapes

For each measure, we computed the distance of every morphospace network to the empirical consensus connectome (see Section 4.1). Aggregating the replicates per (*η, γ*) combination yields one distance landscape per measure (see Figure 1B). In this landscape, each pixel encodes the mean distance of the 10 networks at that parameter combination to the empirical connectome; in the visualization darker regions indicate parameter combinations producing more brain-like networks. Two representative networks sampled from different landscape regions illustrate the diversity of topologies the model can produce: the first appears nearly random, the second shares more features with the empirical connectome, with high intrahemispheric connectivity and non-random structure. The landscapes of all 16 distance measures can be found in Figure 3A.

**Figure 3:**
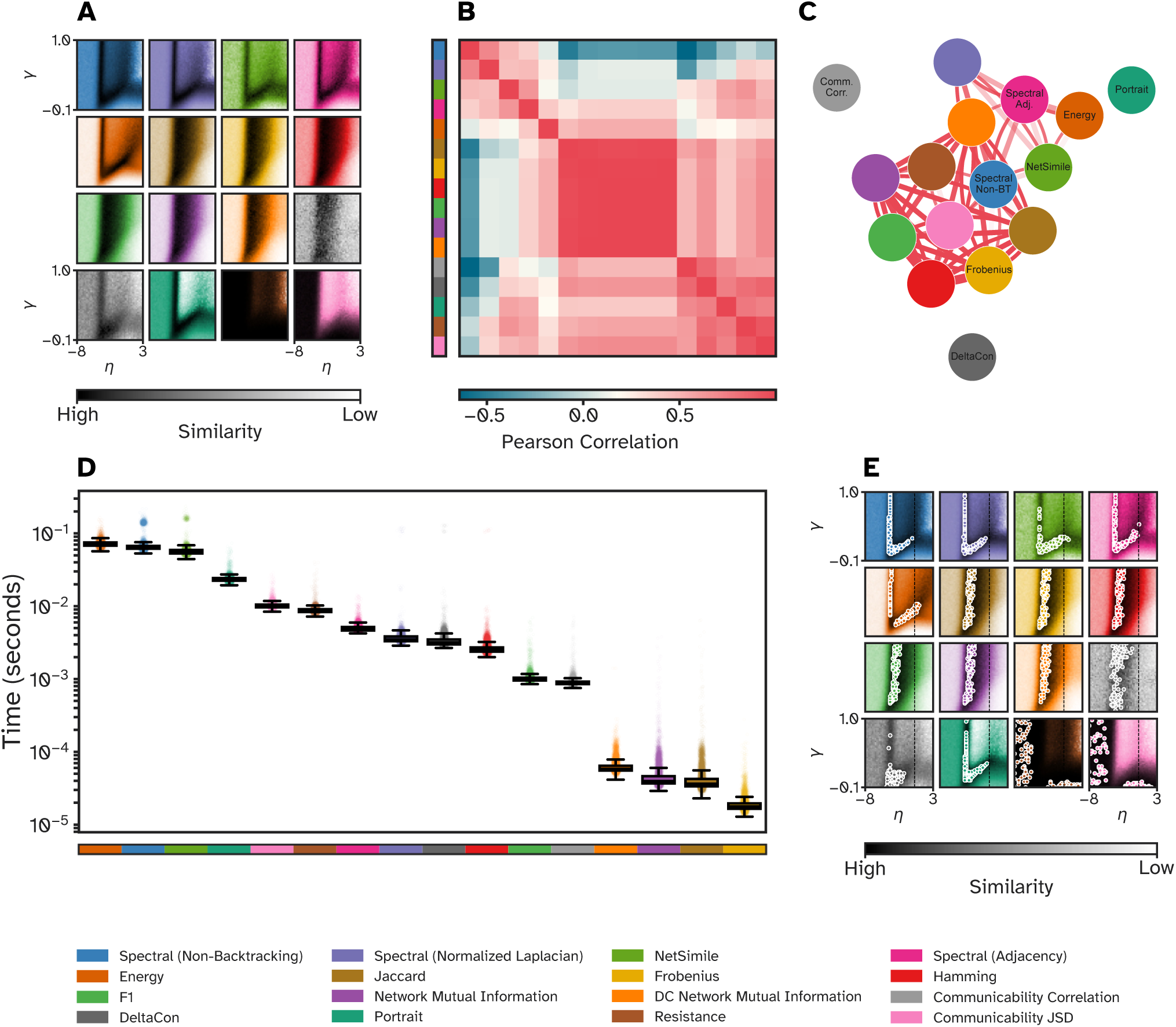
Comprehensive comparison of 16 network distance measures. **A**: Distance landscapes for all 16 distance measures evaluated on the same 25,000 generated networks, with *η* on the horizontal and *γ* on the vertical axis; darker colours indicate higher similarity to the empirical consensus connectome. Three landscape shapes recur: a ’V-shaped’ pattern (e.g. portrait divergence, KS-based energy, spectral (adjacency)), a near-vertical band (e.g. Frobenius, Hamming, NMI), and an ’L-shaped’ pattern (spectral (non-backtracking), spectral (normalized Laplacian)). **B**: Pearson correlation matrix between all landscapes, ordered by hierarchical clustering. Every measure is oriented as a distance first, so a positive entry always denotes agreement. **C**: Inter-measure correlations represented as network graph. An edge is drawn for the 30 strongest pairwise correlations by magnitude. Edge thickness and opacity visualize the strength of the correlation and edge colour its sign (red for positive - all edges of the 30% strongest correlations by magnitude are positive). **D**: Computational runtime per comparison across all 16 measures (log scale), from ∼ 10^−5^ seconds (Frobenius, Hamming) to ∼ 10^−1^ seconds (KS-based energy, spectral (non-backtracking)). Each point is one comparison; the box marks the median and interquartile range. **E**: Locations of the 100 best-fitting parameter combinations (lowest mean distance to the consensus) for each measure in *η*-*γ* parameter space, on the same axes as **A**. The vertical dashed line at *η* = 0 marks the boundary beyond which long-range connections are encouraged rather than penalized.

### Agreement

How far do different measures agree, and where do they differ qualitatively? The landscapes show clear overlaps and differences between measures, see Figure 3A. To elaborate, while most measures show the same V-shaped structure (e.g. portrait divergence, KS-based energy, spectral (adjacency)), a few result in a thicker vertical line (e.g. Frobenius, NMI, and DC-NMI), and some others tend to have an L-shaped structure (e.g. spectral (normalized Laplacian and non-backtracking) and communicability JSD). In the landscapes, the clear V-shaped pattern reflects the competing influences of the two parameters, and the space around the V correspond to the two ways this balance can fail: when *η* is too negative (here: smaller than approximately −4), the distance term progressively stops discriminating between node pairs: The term 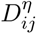 falls below the small constant that the implementation adds to keep wiring probabilities strictly positive^36^ for an increasing share of pairs, until by *η* = *−*6 almost none of them are distinguished, the wiring distribution flattens, and connections are drawn close to uniformly at random. The resulting networks are not short-wired but randomly wired. For instance, at *η* = *−*8 their mean connection length is 68.9 mm, against 80.0 mm for random wiring on the same nodes and 48.7 mm in the consensus connectome. Thus, these networks generated with *η* = *−*8 are quite different to the consensus connectome, regardless of the *γ* value (see Section S5). Networks generated with *η* between −4 and −3 are evaluated as more brain-like, and the closest match to the consensus connectome sits just above *γ* = 0. Along this edge the landscape is almost flat in *γ*, so networks generated with higher *γ* values stay nearly as similar and the low-distance region continues upward as the near-vertical left arm of the V. When *η* is positive, long connections are actively incentivized; since the empirical connectome is dominated by short-range connections - long ones cost more metabolic energy to generate and maintain^37^ - such networks are by default less similar to it. If *γ* is raised so that the (*η, γ*) combination lies on the right arm of the V, the homophily term restores enough of the empirical topology to partly compensate. Two descriptive landscapes over the same grid - the fraction of connections longer than 90 mm and the fraction of interhemispheric edges of the generated networks - support this reading: the parameter combinations that reproduce the empirical values of both properties follow the right arm of the V (see Section S5).

We correlated the landscapes of all 16 measures with each other to show which measures extract a shared intuitiion about network similarity (Figure 3B and C). Agreement is common but not perfect (the pairwise correlations run from *r* = 1.00 down to *r* = *−*0.64) and correlation does not sort the measures into a few clean groups. What the correlation matrix shows instead is one tight block with looser attachments to it.

The tight block is made up of Frobenius, Hamming, Jaccard, F1, NMI, and DC-NMI, that order the morphospace almost identically (*r ≥* 0.973 for every pair of them).

The measures that read global structure, such as spectral (non-backtracking), spectral (normalized Laplacian), spectral (adjacency), NetSimile, and KS-based energy, agree among themselves only moderately (*r* between 0.61 and 0.93) and connect to that block through weaker correlations (e.g. spectral (non-backtracking) with Frobenius, *r* = 0.69).

Communicability JSD and resistance distance sit outside of this tight block. They correlate with each other at *r* = 0.71 and are negatively correlated with KS-based energy (*r* = *−*0.64 and *r* = *−*0.39), i.e. they are ordering the morphospace partially in the opposite direction than the other measures.

Three measures attach nowhere: portrait divergence, DeltaCon, and communicability correlation carry no edge in Figure 3C, because none of their correlations reaches the cut of the 30 strongest pairwise correlations by magnitude (their strongest correlation with any other measure is *r* = 0.65, *r* = 0.64, and *r* = 0.53, respectively).

Portrait divergence^16^ perfectly shows such an intermediate position: the measure takes path-length distributions at all structural scales into account, and thus correlates with other global-structure measures (spectral (normalized Laplacian) *r* = 0.65, DeltaCon *r* = 0.64, NetSimile *r* = 0.58). It is, however, close to uncorrelated with the edge-identity measures, which reward matching specific edge positions rather than global path topology: *r* = 0.09 with Hamming and *r* = 0.02 with DC-NMI.

In conclusion, one network can appear similar to a reference network when utilizing one measure and distant when utilizing another measure. To analyze the similarity and differences between the measures more rigorously, we performed a Principal Component Analysis (PCA) on the landscapes of the eight measures selected based on the filtering procedure described in the next paragraph (see Section 4.3). It reveals a dominant component: PC1 explains 55.1% of the variance and captures the consensus across all measures indicating that - despite distinct mathematical formulations - they converge on a common notion of network similarity. PC2 (16.1%) is the axis of differentiation: measures with high PC2 loading are sensitive to fine-grained, local structure in the morphospace, while measures dominated by PC1 reflect coarser, global differences. A full analysis is provided in Section S4.

### Biological plausibility

Do distance measures select biologically plausible networks? We identified per measure the 100 grid cells whose networks have, on average, the lowest distance to the empirical connectome (Figure 3E). Two kinds of selection recur throughout this work and are *not* interchangeable: a measure’s *n best-fitting (parameter) combinations* are the *n* grid cells with the lowest distance averaged over their 10 replicates, whereas its *n best-fitting networks* are the *n* individual generated networks with the lowest distance of their own. The spatial distribution of the 100 best-fitting combinations varies considerably: for resistance distance and communicability JSD the locations are widely scattered, while for most other measures they cluster tightly around a point (e.g. DeltaCon, the desirable behaviour) or along the vertical line at *η ≈ −*4 (e.g. Frobenius, this spread across all *γ* values implies insensitivity to homophily’s contribution). The vertical dashed line at *η* = 0 marks a biological plausibility boundary: networks with *η >* 0 are actively incentivized to form long-range connections, producing wiring patterns that are not consistent with the metabolic constraints of real brains. Five measures place best-fitting combinations in this implausible region: resistance distance places 27 of its 100 at *η >* 0, communicability JSD and KS-based energy 18 each, NetSimile 4, and spectral (adjacency) 1; the remaining eleven measures place none.

We then used two criteria, biological plausibility and redundancy, to reduce the set of analyzed distance measures from 16 to 8: On grounds of biological plausibility, resistance distance and communicability JSD were excluded, both of which place implausible combinations among their 20 best. Among the remaining measures, Frobenius, Hamming, Jaccard, F1, NMI, and DC-NMI carry a pairwise correlation of at least 0.973 with one another (see Figure 3B and C) and thus, we only retained Frobenius as it is the measure with the lowest computation time of the group (see Figure 3D; more information on computational efficiency will be given in the next paragraph). Spectral (normalized Laplacian) was also excluded due to redundancy: it correlates at 0.926 with spectral (non-backtracking). This is a looser redundancy than the one binding the Frobenius group, and it is the one selection step in this filter that rests on a judgement rather than on a threshold. The subsequent analyses therefore use eight measures: Frobenius, DeltaCon, spectral (non-backtracking), spectral (adjacency), communicability correlation, portrait divergence, NetSimile, and KS-based energy. To increase intuition for these measures, Figures S1-S6 in Section S1 and Figure 2 in the main text provide a visualization of the mechanics of the selected distance measures by displaying all intermediate steps in the calculation of each distance measure, illustrated on one example network pair.

How consistently does each measure locate a region of parameter? And do the networks it selects resemble real connectomes in their fundamental structural properties? To answer the first question, we computed the spread of a measure’s 20 best-fitting parameter *combinations* in parameter space (Figure 4A). To answer the second question, we computed the structural properties of its 20 best-fitting *networks* (Figure 4B and C). Starting with the first question about the spread of each measure’s best-fitting parameter combinations: The total variation in the locations of those 20 combinations - the summed variance of *η* and *γ* across them, each normalised to the extent of the morphospace (Figure 4A) - reveals substantial differences in precision: DeltaCon (0.003) and KS-based energy (0.004) identify a small, consistent region of parameter space, followed by NetSimile (0.010), portrait divergence (0.014), spectral (adjacency) (0.023), and spectral (non-backtracking) (0.049), while Frobenius (0.078) and communicability correlation (0.109) spread over an order of magnitude more of the morphospace.

**Figure 4:**
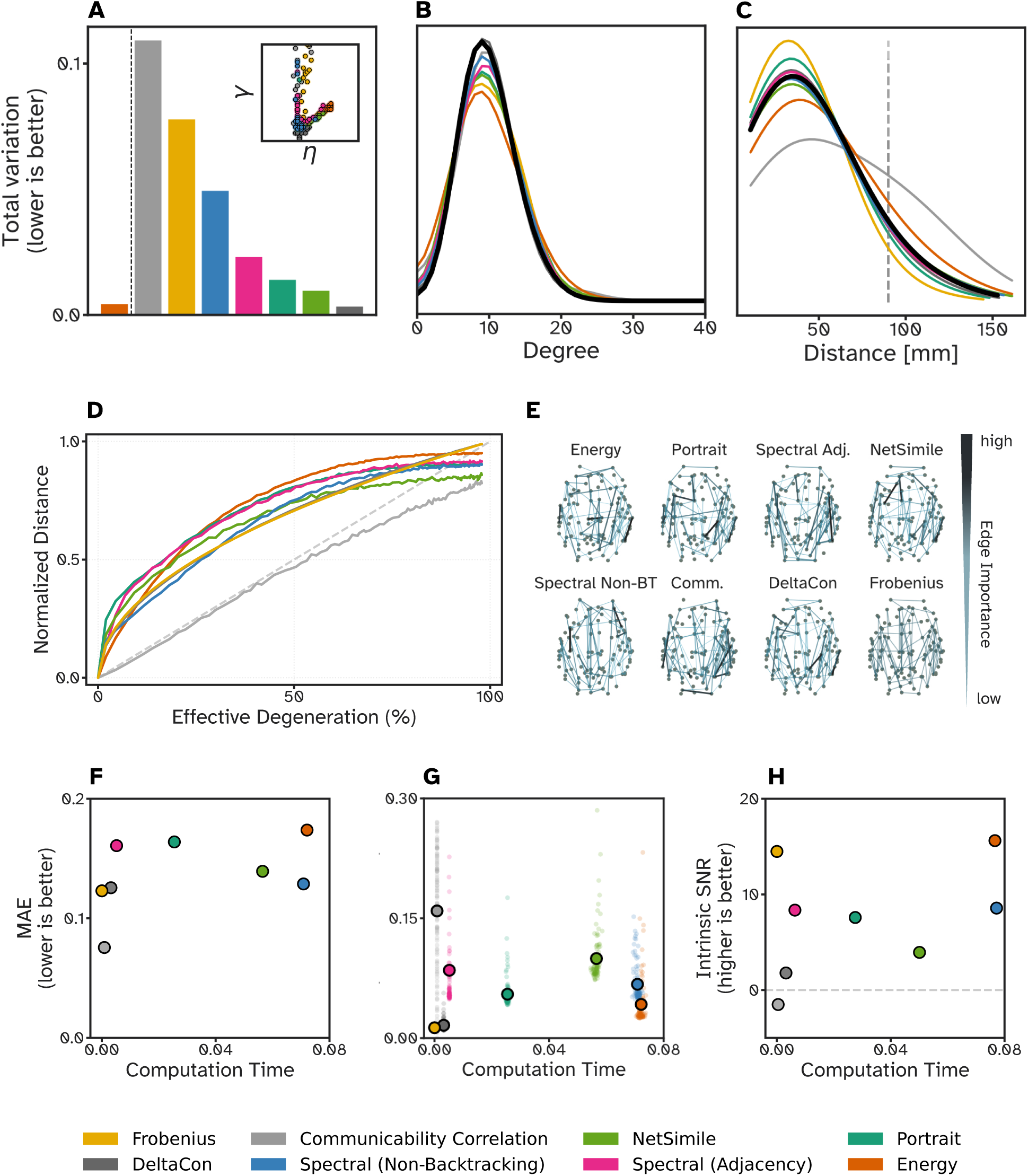
Precision of the selected parameter regions and reaction to perturbations. **A, inset**: Locations of the 20 best-fitting parameter combinations per measure. **A, main**: Their total variation, var(*η*) + var(*γ*) over the 20 cells, normalised to the extent of the morphospace. **B and C**: Degree and edge distance distributions of each measure’s 20 best-fitting networks (coloured) and of the 100 individual empirical connectomes (black). The vertical line in **C** marks the 90 mm threshold for long-range connections used by Oldham et al.^13^. **D**: Response of each measure to progressive degree-preserving rewiring of the consensus, against the effective degeneration, i.e. the fraction of edges that actually differ from it (see Section 4.10). Both axes are min-max normalised within each trajectory and then averaged over the 200 trajectories; a trajectory’s largest value need not fall at its last level, so the averaged curves end short of the top right corner. The dashed diagonal marks a linear response. **E**: Edge-importance maps. Nodes are drawn at their anatomical locations (projected from above) and the top 20% most important connections shown; colour and width encode the distance increase caused by removing that connection (wide and black: large; thin and blue: small). **F**: Computational cost against deviation from the linear response in **D**, measured as the mean absolute error (MAE) between the curve and the diagonal. **G**: Computational cost against measurement stability, the coefficient of variation across the 200 trajectories at each perturbation level (see Section 4.10); the mean per measure is the larger point. **H**: Intrinsic signal-to-noise ratio (iSNR) of each landscape, the ratio of signal variance to replicate noise variance (see Section 4.9).

Both the total variation and the analysis of the selected networks provide complementary insight: the total variation says how consistently a measure points at one region of parameter space (presision), but a measure can be perfectly consistent about a region that produces unrealistic networks. The answer to the second question therefore complements the plausibility axis begun above, asking whether the networks a measure ranks closest carry biologically plausible degree distributions (Figure 4B) and connection distance distributions (Figure 4C). The empirical reference in both panels is the set of 100 individual connectomes, which fixes the range a generated network would have to fall in. The degree distributions barely separate the measures: all eight sit close to the empirical distribution (black curve; the 100 individual connectomes). The nodal degree of the individual connectomes have a standard deviation of 3.49 ± 0.20 across the 100 nodes. DeltaCon (standard deviation of nodal degree of 3.54) and spectral (non-backtracking) (3.65) fall inside that empirical spread, communicability correlation (3.79), spectral (adjacency) (3.81), portrait divergence (3.87), Frobenius (3.90) and NetSimile (4.02) sit just outside it, and KS-based energy is clearly the broadest (4.55).

The connection distance distributions separate them more sharply and in both directions. Real connectomes average 42.1 ± 1.1 mm per connection and place 6.6 ± 1.2% of their connections beyond 90 mm - the threshold for long-range connections utilized in^13^. Across the 100 individuals these statistics span 39.5 to 44.8 mm and 3.4 to 9.3%, and four measures select networks that fall inside both spans: NetSimile (41.8 mm, 7.4%), spectral (non-backtracking) (41.3 mm, 7.2%), spectral (adjacency) (40.9 mm, 6.2%) and DeltaCon (39.7 mm, 4.6%). Portrait divergence (38.7 mm, 3.4%) sits at the short edge of that range and Frobenius (35.4 mm, 2.0%) well below it, retaining less than a third of the empirical share of long connections. The two deviations in the other direction are larger: KS-based energy selects networks averaging 47.6 mm with 10.5% of connections beyond 90 mm, and communicability correlation 60.8 mm with 25.5%, nearly four times the empirical share.

KS-based energy’s overshoot is (at least partially) explained by a potential limitation of this experiment. The measures are fitted against is the *group* consensus (not the 100 individual connectomes). However, the group consensus is itself longer-wired than any individual brain (48.7 mm, 9.7% beyond 90 mm, against a maximum of 44.8 mm across the 100 individuals). As the KS-based energy’s objective contains a specific edge-length term, it might be that the measure faithfully reproduce the consensus’s distribution, instead of reproducing the generally observed degree and distance distributions of individual (biological) brain networks.

Both criteria just applied are distributional. However, even if a network can reproduce the degree distribution of the reference exactly, it can still put every hub in the wrong region. Thus, evaluating the measures on a topographic criterion is interesting, and we therefore tested whether the criterion can be sharpened into a topographic criterion. Looking at topography is only possible because the generated and empirical networks are built on the same Schaefer-100 parcellation and are thus node-aligned. The most direct form of a topographic criterion is to correlate nodal degree across the 100 nodes between each generated network and the empirical consensus (see Section 4.5; full analysis in Section S6). This sharpening fails, but it fails due to the generative model rather than due to the measures: No generated network comes close to the empirical hub topography - across all 2,500 parameter combinations the best agreement is *r* = 0.21 (and only *r* = 0.09 for the best individual generated network) - whereas single empirical connectomes correlate with the same consensus at *r* = 0.68 *±* 0.06, the weakest of them still at *r* = 0.41. A distance measure can only select among these generated networks, so no selection it makes can reach the empirical range. Instead of the empirical hub topography, the generated networks have one that is determined by geometry: nodal degree tracks spatial centrality, whereas the empirical degree map does not (see Figure S14). In conclusion, this direct form of a topographic criterion thus measures the generative model rather than the measure applied to it. Thus, we do not use it as a plausibility axis.

### Computational efficiency

How do the measures compare in computational effort? Runtime varies by approximately four orders of magnitude across measures (Figure 3D). KS-based energy is the most expensive (71.72 ± 6.31 ms per comparison), followed by spectral (non-backtracking; 67.48 ± 16.27 ms) and NetSimile (57.23 ± 10.63 ms); Frobenius is the fastest (0.019 ± 0.006 ms). Fitting one morphospace of 25,000 networks to one connectome therefore costs 29.9 minutes more with energy than with Frobenius - scaling to 13.52 days if one morphospace is refitted per subject across the 652-subject HCP-D dataset^38^. Runtime is therefore a practical constraint: for large datasets, faster measures may be necessary even at the cost of other desirable properties.

### Accuracy

Can distance measures recover known generative parameters? All analyses described above lack a true ground truth: as there does not exist one unique parameter combination that gave rise to each empirical connectome, we can only assess *precision*, but not *accuracy*. We therefore conducted a synthetic parameter recovery experiment in which 100 ground-truth parameter combinations were drawn uniformly from the whole morphospace and one test network was generated at each of them, and each distance measure was used to identify the closest location on a dedicated 10 × 10 recovery grid covering the whole morphospace (*η ∈* [*−*8, 3], *γ ∈* [*−*0.1, 1], so that one grid step is 1.22 in *η* and 0.12 in *γ*) (Figure 5A, Figure 5E left; see Section 4.7; full results in Figure S16 in the Supplementary Material). The recovery error is expressed in grid steps, defined as the Euclidean distance in grid-index space between the true and predicted cell. The maximum possible error on a 10 × 10 grid is approximately 12.73 steps. DeltaCon achieves the most accurate recovery (2.736 grid steps), followed by F1 and Jaccard (both 3.177) and KS-based energy (3.234). The measures performing the most poorly are DC-NMI (4.828 grid steps), spectral (normalized Laplacian) (4.206 grid steps), spectral (non-backtracking) (4.086 grid steps), and portrait divergence (3.772 grid steps). No measure exceeds the 4.96 grid steps expected of a measure recovering a uniformly random cell of this grid (see Section 4.7).

**Figure 5:**
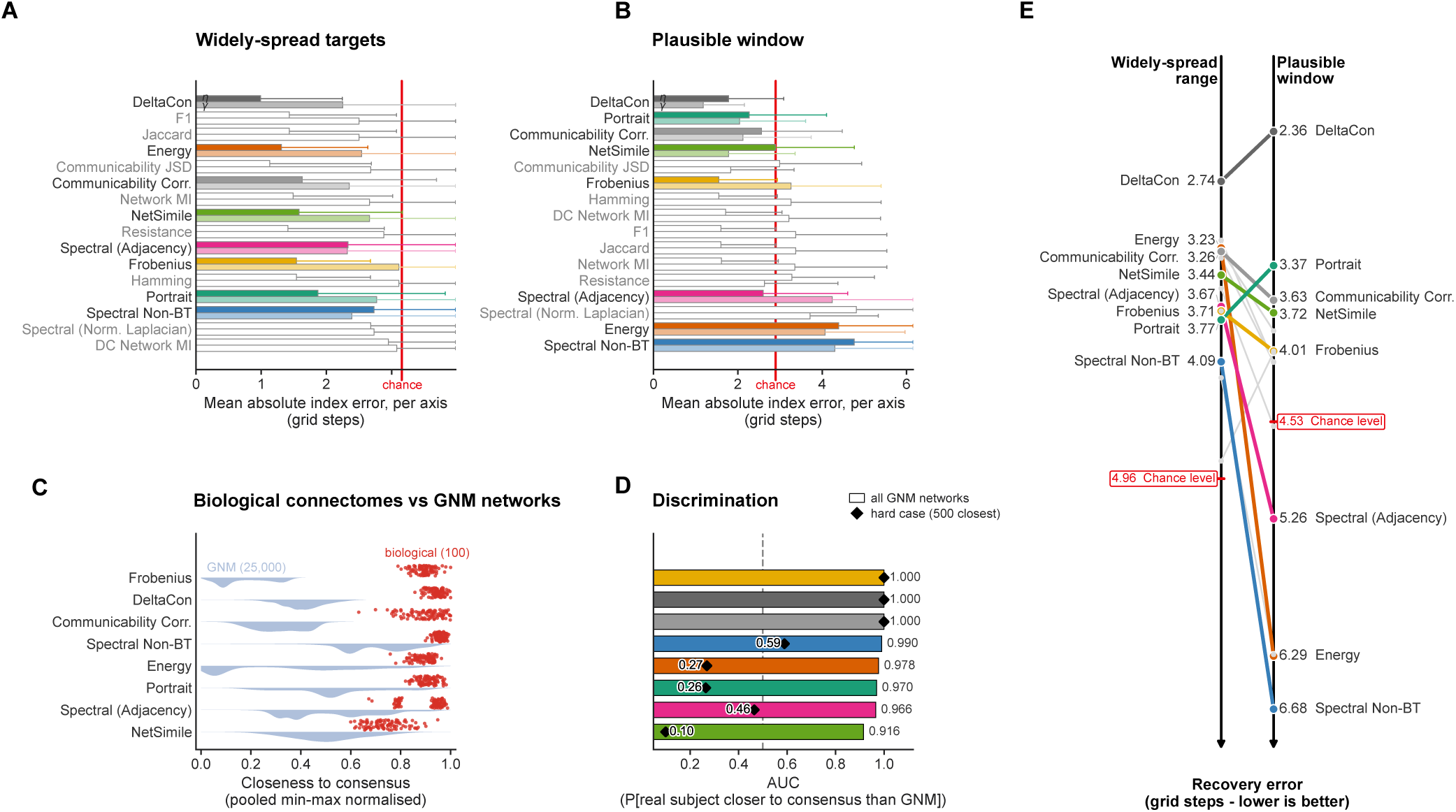
Ground-truth parameter recovery and real-versus-synthetic discrimination. **A**: Recovery error on 100 ground-truth parameter combinations drawn uniformly from the whole morphospace (*η* ∈ [−8, 3], *γ* ∈ [−0.1, 1]; one test network each). Bars give the mean absolute index error along *η* (dark) and along *γ* (light) on the 10 × 10 recovery grid, averaged over the test networks; measures are ordered by the mean Euclidean distance in grid-index space, which is also used in panel **E**. Whiskers are one-sided: each starts at the mean and extends outwards by one standard deviation of the per-network error. The red line marks the per-axis error of a measure recovering a uniformly random cell (3.16 grid steps here, 2.90 in **B**). **B**: The same read-out for the 100 ground-truth networks drawn from the plausible window (*η* ∈ [−3.96, −1.49], *γ* ∈ [0.08, 0.30]), recovered on a dedicated 10 × 10 grid covering that window. **C**: Closeness-to-consensus distributions for the eight selected measures. Light blue violins show the 25,000 GNM networks, scored against the full consensus, and red points the 100 real HCP subjects, each scored against a consensus rebuilt from the other 99 subjects so that no subject contributes to its own reference. Closeness is pooled min-max normalised per measure, and higher values indicate networks closer to the consensus. **D**: Discrimination ranking. Bars show the AUC computed against all 25,000 GNM networks, i.e. the probability that a random real subject is closer to the consensus than a random GNM network. Black diamonds show the AUC computed against each measure’s 500 closest GNMs. The gray line marks chance (0.5). **E**: Mean grid-step error of each of the 16 measures on the widely-spread targets (left) and inside the plausible window (right), with the two values given per measure and a line connecting the same measure across the two experiments. The eight selected measures are drawn in their colour and the remaining eight in grey; the red markers give the chance level of each experiment (4.96 grid steps on the widely-spread targets, 4.53 inside the window). Vertical position encodes the error.

DeltaCon is the measure that stands apart: the twelve measures ranked behind it, from F1 to portrait divergence, span barely six tenths of a grid step, which is far less than the spread of the per-network errors within any one of them (Figure 5A), so they are best read as one middle group rather than as a ranking.

In Figure 5A it is interesting to observe that almost every measure recovers the value for *η* better than the value for *γ*. The two spectral measures that were selected for the benchmark are the exceptions, recovering *γ* at approximately as well as or better than *η*.

These targets, however, span the entire morphospace, whereas empirical work operates inside a much narrower window of parameter space. We therefore repeated the experiment for the same 16 measures. Again, we drew the 100 ground-truth networks, but this time from the *plausible* window alone (*η ∈* [*−*3.96*, −*1.49], *γ ∈* [0.08, 0.30], that we derived from the pooled interquartile range of the locations of the 100 best-fitting networks of each of the eight selected measures against the empirical consensus; see Section 4.7). Then, parameter recovery was performed for each measure on a dedicated 10 × 10 grid over *η ∈* [*−*4.31*, −*1.14] and *γ ∈* [0.05, 0.34] (again slightly wider than the window generating ground-truth networks, so that recovery is not truncated at its edges) and scoring recovery in grid steps as above. One grid step is 0.35 in *η* and 0.03 in *γ* here, against 1.22 and 0.12 on the wide grid, so a grid step is a much finer distinction in this experiment and the two sets of scores are each read against their own chance level rather than against each other (Figure 5E, with the per-axis decomposition in panel 5B; see Section 4.7 and Section S8). Recovery inside this window is naturally more difficult for any measure.

The best score is 2.36 grid steps (DeltaCon), followed by portrait divergence (3.37), communicability correlation (3.63), and NetSimile (3.72). Five of the 16 measures score at or beyond the 4.53 grid steps expected of a measure recovering a random cell in this window (see Section 4.7): resistance (4.57), spectral (adjacency) (5.26), spectral (normalized Laplacian) (6.28), KS-based energy (6.29), and spectral (non-backtracking) (6.68). The two spectral measures that were selected for the benchmark are also the ones whose recovered parameters were nearly constant (both returned six distinct cells as recovered parameters for the 100 ground-truth networks). The ranking by grid steps barely survives the change of window size(Figure 5E). DeltaCon is the measure that leads in both regimes.

Do measures rank real empirical connectomes as more similar to the consensus network than synthetic GNM networks? As a model-free addition to the parameter recovery analysis, we tested whether each measure scores each of the 100 real individual connectomes as closer to the empirical consensus than *all 25,000 synthetic GNM networks* (Figures 5C and 5D; see Section 4.8). Each subject was scored against a consensus rebuilt from the other 99 subjects, so that no subject contributes to its own reference. Against the full GNM population, every selected measure separated real from synthetic networks almost perfectly (AUC of at least 0.916 for all eight, see scatter vs. distribution plot in Figure 5C). This result confirms that no measure is grossly inverted. A second, harder test separates the measures far more sharply (Figure 5D). In our prediction, empirical connectomes carry structure that the generative model does not reproduce, so a measure that is sensitive to that structural element should keep every real subject closer to the consensus than the generated networks - even for the generated networks that it itself ranks as best-fitting. We therefore repeated the comparison against each measure’s *own 500 best-fitting networks* rather than against the full GNM population. Frobenius, DeltaCon, and communicability correlation retain an AUC of 1.00 (i.e. real subjects stay closer to the consensus network than every one of those 500 networks), while the AUC for spectral (non-backtracking) and spectral (adjacency) falls to 0.59 and 0.47, respectively. KS-based energy (0.27), portrait divergence (0.27), and NetSimile (0.10) drop well below chance, scoring their 500 best-fitting networks as more similar to the consensus network (i.e. as more brain-like) than the median real connectome. To rephrase: only three of the eight measures keep every real subject ahead of their own best generated networks, which is the expected behaviour. A measure can recognize that real connectomes are closer to the consensus than a huge set of unrelated networks, yet still mistake its own best-fitting networks for a real brain.

### Sensitivity and robustness

How robust and sensitive are distance measures to network perturbations? We subjected the consensus network to progressive random rewiring (101 levels, from no swap to one attempted swap per edge) while preserving its degree distribution, and report each measure’s response against the effective degeneration rather than against the attempted swaps (Figure 4D; see Section 4.10). The distance at each level was averaged over 200 independent rewiring trajectories. An ideal measure should exhibit - for most use-cases - monotonic, approximately linear increases with perturbation magnitude (i.e., the absolute difference between the adjacency matrix and the rewired adjacency matrix), demonstrating proportional sensitivity to both small and large structural changes.

All measures rise monotonically, and seven of the eight run above the diagonal: they respond more strongly to the first edges that change than to the last ones. Portrait divergence, spectral (adjacency), and NetSimile have the steepest slopes at low effective degeneration, reaching a third of their range once a tenth of the edges differ. KS-based energy and spectral (non-backtracking) start shallower, but KS-based energy then rises above every other measure from roughly a quarter of the edges onwards and stays there until the final stretch, where Frobenius and DeltaCon catch up with it. Frobenius and DeltaCon follow nearly identical trajectories throughout. Communicability correlation is the exception in two ways: first, it responds most gradually by tracking the diagonal more closely than any other measure. Second, it is the only measure that runs *below* the diagonal, meaning it under-reports edge changes rather than over-reporting them. Communicability correlation, Frobenius, and DeltaCon are also the three measures that saturate last, still gaining about a seventh of their range over the top quarter of the perturbation axis while the other five gain less than a twentieth.

That later saturation allows Frobenius, DeltaCon, and communicability correlation to remain sensitive to structural changes even when networks are already substantially perturbed. Together, these results highlight a key consideration when selecting a measure: sensitivity is not uniform across the range of structural differences. Measures with a steep initial slope are best suited for comparing highly similar networks (e.g. when modeling individual differences), but may overreact when comparing broadly different networks (e.g. across species).

The analysis above perturbs the networks being compared. A separate question is what happens to the best-fitting (*η, γ*) combination, if a few of the edges of a network had been placed elsewhere. Would each measure still select the same best-fitting parameter combination? To answer this, we rewired *s* edges of the consensus while preserving its degree sequence, density, and connectivity, re-recovered the best-fitting (*η, γ*) combination at each noise level, and measured the drift from the unperturbed best-fitting combination in grid steps (see Section 4.12, Section S9, and Figure S17). Tolerance to this kind of noise spans close to two orders of magnitude: communicability correlation holds its best-fitting combination until 26% of the edges have effectively moved, DeltaCon until 6%, Frobenius and NetSimile until 3%, and the two spectral measures until 1.5%. KS-based energy and portrait divergence give way at 0.7%, roughly two swaps of the 495 edges. Portrait divergence is in practice the least tolerant of the eight: it is the only measure whose best-fitting combination already moves at the smallest perturbation tested. For it, a single swap displaced the best-fitting combination by approximately 1.5 edges (median drift 1.1 grid steps), while the other seven hold their best-fitting combination.

Next, we measured the distance change caused by removing each connection of the consensus network individually (Figure 4E; see Section 4.11), to see which edges each measure weighs most heavily. Edges whose removal produces the largest distance increase are considered most important by a given measure. We visualized the top 20% of connections ranked by importance for each measure, with edge color and width encoding importance (wide and black: high; thin and blue: low). Energy, spectral (non-backtracking), and DeltaCon identify short-range edges as disproportionately important, while spectral (adjacency) and communicability correlation prioritize longer connections. Notably, Frobenius treats all edge removals equally, reflecting its connection-by-connection approach. Together, these results leave two trade-offs to weigh against the experimental context: a more sensitive measure resolves small differences but saturates earlier, and the sensitivity it adds is often bought with a large increase in computation time.

What trade-offs exist between measurement quality and computational cost? As shown in Figure 3D, different measures require vastly different computational resources, motivating us to evaluate performance relative to cost. To do so, we assessed three complementary quality indicators, see Figures 4F-H.

First, we quantified response linearity as the mean absolute error (MAE) between each measure’s rewiring response curve and the effective degeneration it should track, i.e. the diagonal of Figure 4D (Figure 4F; see Section 4.10). Each measure contributes a single point, the MAE of its mean response curve over the 200 rewiring trajectories (101 rewiring levels from 0-100%). Communicability correlation achieved the lowest MAE (0.076), substantially outperforming all other measures (which all lie above 0.12). The next-best performers were Frobenius (0.123) and DeltaCon (0.126). In addition to its computational cost, KS-based energy exhibited the highest deviation from linearity (MAE = 0.174). This is not a problem per se, but could be one if the research question does not require the sensitivity that the measure offers.

Second, we quantified measurement stability using the coefficient of variation (CV) across the 200 rewiring trajectories at each perturbation level (Figure 4G). Note that the y-axis excludes the eight points with a CV above 0.3: three for spectral (adjacency), at CVs of 0.4307, 0.4919, and 0.9969, and five for communicability correlation, at CVs between 0.3041 and 0.5061. All eight lie at computation times below 0.006 seconds. Frobenius (CV = 0.0130) and DeltaCon (CV = 0.0161) demonstrated the highest stability, followed by energy (0.0422), portrait divergence (0.0549), and spectral (non-backtracking; 0.0674). Communicability correlation showed substantially higher noise (CV = 0.1536).

Lastly, we computed the intrinsic signal-to-noise ratio (iSNR) across the entire parameter landscape (Figure 4H), quantifying each measure’s ability to distinguish meaningful network variation from noise (see Section 4.9). KS-based energy (15.62) and Frobenius (14.49) reach the highest values and are not separable from one another, followed by spectral (non-backtracking) (8.57), spectral (adjacency) (8.36), and portrait divergence (7.59). NetSimile (3.94) and DeltaCon (1.79) keep only a small margin of landscape structure over their replicate noise, and communicability correlation is the one measure with a negative iSNR (−1.51): the spread between its ten replicates at a given parameter combination is larger than the spread of its landscape across combinations. The ratio is scale-invariant, so measures whose raw values live on different numeric scales are directly comparable here.

Together, these analyses reveal that *no single measure excels across all quality dimensions*, necessitating careful consideration of specific application requirements when selecting distance measures for generative network model evaluation. We translate these trade-offs into concrete, use-case-specific recommendations in the Discussion.

### Comparison across axes

To summarize the practical trade-offs, we condensed each of the five evaluation axes into one spoke of a radar plot per measure (Figure 1D; see Section 4.13). Every axis is represented by the single read-out from the described experiments that reduces most cleanly to one number per measure. The scores are ordinal and are meant as a qualitative summary of trade-offs. This visualization highlights the compromises that are inherent to each measure. For instance:

Frobenius is the cheapest by a wide margin, but it clears the chance level for recovery only narrowly. Additionally, the consistency of the region it selects is lower than for all other measures except communicability correlation.

KS-based energy, on the other hand, is the slowest of the eight, and is, inside the plausible window, less accurate than picking a cell at random (6.288 grid steps against a chance level of 4.53). Its value, however, is built from properties the field already interprets (i.e. degree, clustering coefficient, betweenness centrality, and edge length), so a poor fit can be traced back to the property responsible for it.

Portrait divergence also requires long runtimes, but achieves the second best accuracy inside the plausible window. It has the least tolerance to a perturbed reference.

DeltaCon has the highest values across the five evaluation axes: It ranks first on the consistency of its selected region, first on accuracy, and second on perturbation robustness. While doing that, its runtime is roughly twenty times shorter than energy’s.

## 3 DISCUSSION

Methods for comparing networks in general have been benchmarked before, and prior work in the field of network neuroscience has compiled the measures in circulation and called for a systematic comparison on simulated data^22^, which is exactly the comparison we set out to perform in this study. We evaluated 16 measures on 25,000 synthetic networks grown with a generative network modelling (GNM) framework. From the measures, a subset was filtered, and the remaining 8 measures were scored on five axes: agreement, biological plausibility, computational efficiency, sensitivity and robustness, and accuracy. No measure leads on more than two of the five, and the measure the field has standardised on leads on none.

Most measures agree on the overall shape of the landscape (i.e. converge on the same V-shaped landscape (Figure 3A)), and one principal component captures 55.1% of the variance across the eight selected measures. As the general shape of the landscape is a consequence of the generative model (at strongly negative *η* the model wires randomly rather than short, at *η >* 0 the model is rewarded for the long connections that are scarce in the empirical connectome), and any measure with a working sense of structure registers both boundaries, the high agreement observed is expected. The informative component is PC2 (16.1%), which separates measures sensitive to local structure from those sensitive to global structure. Generally, agreement is the one axis on which a high score certifies a shared sense of intuition of network similarity to other measures, rather than correctness.

Agreement on the landscape does not carry over to the networks and which parameter-combinations each measure sees as best-fit to the consensus connectome. Resistance distance and communicability JSD place best-fitting combinations where long-range connections are rewarded and thus no metabolically constrained brain could follow, and are excluded on that basis. Among the eight that remain, the 20 closest generated networks separate sharply and in both directions: NetSimile, the two spectral measures and DeltaCon fall inside the range spanned by the 100 individual connectomes on both connection-length statistics, while Frobenius selects networks far too short-wired (35.4 mm against 42.1 mm across individuals), and communicability correlation (60.8 mm) and KS-based energy (47.6 mm) too long. The overshoot of KS-based energy might be faithful: the group consensus it is fitted to is longer-wired than any the individual connectomes that teh consensus was built from (48.7 mm), and the KS-based energy might detect this due to edge-length term in its objective. Generally, minimizing the distance value between a synthetic and the real network is not the same as recovering a realistic network.

Parameter recovery is the only test with a known answer. We ran a parameter recovery experiments twice; on targets spanning the full morphospace and on targets in the narrow empirical window. The two rankings barely overlap: portrait divergence rises from 13th to 2nd, while KS-based energy drops from 4th to 15th (below chance). DeltaCon leads both regimes, and our recommendation for it rests on that result.

Cost and stability bound the other axes. Runtime spans approximately four orders of magnitude: fitting one morphospace per subject across 652 HCP-D subjects^38^ takes 13.5 days longer with KS-based energy than with Frobenius. Robustness to reference perturbation spans two orders of magnitude: communicability correlation holds its optimum until 26% of edges have moved; KS-based energy and portrait divergence shift at 0.7%, portrait divergence drifting after a single swap (Section S9).

The five axes do not weigh equally and their value depends on the task. Thus, they are not sensibly combinable into a single score. For instance, a failure on accuracy is a failure against a known answer and might invalidate an entire measure for a certain experimental question, while low agreement means only that a measure differs from its peers. Thus, our measure recommendations depend on the research question or the measure’s purpose at hand (Figure 1D and E). As an all-purpose default we recommend DeltaCon, as it leads both accuracy and precision, performs for all axes above-average (except for the axes of agreement), and runs *∼*20*×* faster than KS-based energy. Its low signal-to-noise ratio and slightly short-wired selections are the measure’s weak points. For landscape visualization, for instance, that signal-to-noise ratio is the deciding factor. For this task, KS-based energy and portrait divergence are the most suitable from all measures that resolve both *η* and *γ* parameters (Frobenius also has a high signal-to-noise ratio, but lacks the ability to resolve *γ*). As KS-based energy is very expensive, portrait divergence is most recommendable; KS-based energy for landscape visualization bears the advantage of creating landscapes that can be compared with prior work. For large-scale application, runtime is critical, and thus Frobenius is the choice. This comes at the price of insensitivity to *γ* and the most short-wired selections of the eight. It may best be used for the fast first screening for a parameter region that produces brain-like networks, before turning to more expensive methods. When spatial constraints are the priority, DeltaCon and KS-based energy are our recommendations, depending if the target are the biologically-plausible reconstruction of individual connectomes, or the best approximation of the group-level connectome, respectively. For parameter optimization DeltaCon leads both parameter recovery regimes.

These recommendations are bound by three limitations: First, the rankings are point estimates with no reported uncertainty, so only the larger gaps are safe to interpret. Second, all results rest on one reference net and one generator, which both have their limitations: The consensus network is longer-wired than the individuals behind it, which might skew results, and the utilized network generation model might not capture the full complexity of brain networks. Third, our plausibility criterion counts connections without locating them, and it cannot easily be sharpened to include where the hubs sit, as the ceiling is set by the network generator rather than the measures (i.e., no generated network reproduces the hub topography of the consensus (*r ≤* 0.21), while every individual connectome does (*r* = 0.68 *±* 0.06). Also, the generated hubs are merely geometric (Section S6)). This is the failure that motivated adding changing brain geometry^23^ and heterochronous growth^24^ to generative models. Adding a similar analysis for a different generative model might provide further insights, if the generative model produces networks with a more realisitc placement of hubs.

To summarize our key findings (Figure 1D and E): in agreement with prior literature, this benchmark observes that distance measures agree on which networks are grossly unlike the utilized reference consensus connectome. However, the measures diverge at declaring which parameter combinations fit best to a reference network, which is, however, often the one step that is the core of a published result. Additionally, minimizing a distance calculated by a measure is not the same as recovering a realistic network. Among the measures that pass our plausibility filter, the best-fitting networks span a third to four times the empirical share of long-range connections, so goodness of fit and biological plausibility have to be reported as two separate criteria. Lastly, no measure serves every purpose: no single measure leads on more than two of the five axes. The current standard in the field, KS-based energy, leads on no axis and performs on the chance level when recovering parameter combinations in the narrow parameter window in which empirical work operates.DeltaCon, while being the closest to an all-purpose default, still has its limitations. Most important to remember is that which measure a study uses decides which networks it selects as most similar to the reference connectome. Thus, the choice of measure should be stated and - ideally - justified; just like it is done for the choice of parcellation.

## 4 MATERIALS AND METHODS

### 4.1 Empirical Consensus Connectome

The structural connectomes used in this analysis were processed as part of^39^. Briefly, diffusion MRI data from 100 unrelated participants in the Human Connectome Project S900 release^40^ were reconstructed using QSDR^41^ in DSI Studio and parcellated according to the Schaefer-100 atlas^42^. Deterministic tractography (modified FACT^43^) yielded 1,000,000 streamlines per subject, from which weighted connectivity matrices were constructed.

These networks were then further processed to a binarized version with a fixed density of 10% of the individual connectomes, using the functions `threshold_network` from `netneurotools`^44^. This function ensures that the network remains connected by utilizing a minimum spanning tree. The individual networks were computed to compare the distance and degree distributions of the empirical networks with the best-fitting networks of each distance measure. Full acquisition and preprocessing details are reported in^39^ and the original HCP documentation^45–47^.

A consensus network was additionally derived across the 100 individual connectomes using the `struct_consensus` function from `netneurotools`. This consensus network is the reference network used for most experiments in this study.

### 4.2 Generative Network Model

We employed a spatially-constrained generative model that constructs networks by optimizing a trade-off between spatial economy (minimizing wiring cost) and topological organization ^11^. The model operates through iterative edge addition, where at each step the probability of forming a connection between nodes *i* and *j* is determined by:

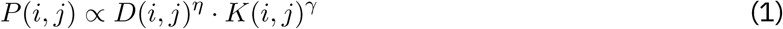

where *D*(*i, j*) represents the Euclidean distance between nodes, *K*(*i, j*) quantifies topological affinity, *η* controls the penalty for spatial distance (negative values discourage long connections), and *γ* modulates the influence of topological similarity, or homophily. The implementation adds a constant of 10*^−^*^8^ to the unnormalised wiring probabilities to keep them strictly positive^36^. This constant sets a floor below which the two terms can no longer be told apart, which is what shapes the landscapes at strongly negative *η* (see Results).

For the utilized homophily-based model, *K*(*i, j*) was created using the ’matching rule’ - a rule that has been shown to create networks that fit best to empirical connectomes (based on the KS-based Energy measure)^12^. The distance matrix *D*(*i, j*) was based on the spatial coordinates of the Schaefer 100 atlas, ensuring that distance penalties reflect realistic brain geometry.

We used binary GNMs, where connections stay unweighted and every matrix entry in the adjacency matrix *A* has either a present (1) or an absent (0) edge.

The generative process begins with a seed network (or no seed network) and iteratively adds edges according to the probability distribution defined above until the total number of connections matches that of the empirical connectome. The seed is retained in the finished network. The seeding network was a Minimal Spanning Tree (MST) generated on the Schaefer 100 atlas distance matrix using the `minimum_spanning_tree` function from `NetworkX` ^48^. This prevents disconnected graphs in the *η*-*γ* parameter space, particularly for high *η* and *γ* values where networks can otherwise fragment into numerous components.

Concretely, each generated network holds the 99 edges of the MST plus 396 edges added by the model, resulting in 495 edges in total. This equals a density of 10%, the same as in the empirical consensus - allowing all comparisons reported here being made at matched density. One consequence of the seeding remains: 41 of the 99 MST edges are absent from the empirical consensus, so those edges are mismatches in every generated network regardless of the (*η, γ*) combination, and no measure can reach zero distance to the consensus anywhere in the morphospace.

We ran the described homophily-based GNM framework for 50 × 50 *η*-*γ*-parameter combinations. The parameters were regularly sampled from *η ∈* [*−*8, 3] and *γ ∈* [*−*0.1, 1.0]. For each of the 2,500 combinations we generated 10 replicate networks to account for stochasticity in the generative process, resulting in 25,000 total synthetic networks.

This replication strategy allows us to distinguish systematic parameter effects from generative noise. Also, the parameter spans include the ranges used in previous literature (sets include for instance *η ∈* [*−*3.5, 0] and *γ ∈* [0, 0.8] in^24^ and in the finer search *η ∈* [*−*3.606, 0.354] and *γ ∈* [0.212, 0.495] in^12^). The parameter ranges were selected to span biologically plausible and implausible regimes. The critical boundary at *η* = 0 separates networks where long connections are penalized (*η <* 0, potentially brain-like) from those where long connections are encouraged (*η >* 0, metabolically implausible). The *γ* range captures varying degrees of homophily, from anti-homophilic connection patterns (*γ <* 0) through neutral (*γ ≈* 0) to strong homophilic influences (*γ* = 1).

The GNM implementation used is from^36^.

### 4.3 Principal Component Analysis of the Landscapes

The correlation matrix shows *which* measures move together but not *what* independent notions of similarity the eight measures carry between them. A PCA answers that by expressing the landscapes in terms of a few shared components, separating the consensus the measures agree on from the axis along which they differ. Each measure contributes one distance per generated network, min-max normalised to [0, 1] across the morphospace. The 25,000 individual generated networks were treated as observations and the measures as variables, the columns were standardised, and a PCA was run on the resulting 25,000 × 8 matrix.

### 4.4 Comparison with Empirical Connectome Properties

Two selections are used in this work and are kept distinct throughout. A measure’s *n best-fitting networks* are the *n* individual generated networks, of the 25,000, with the smallest distance to the consensus network. A measure’s *n best-fitting parameter combinations* are the *n* of the 2,500 (*η, γ*) cells with the smallest distance averaged over their 10 replicates. The two selections do not necessarily coincide, since ten networks share each cell and a cell may be good on average without containing the single closest network.

The precision read-out in Figure 4A uses the 20 best-fitting parameter combinations of each measure: precision is the summed variance of *η* and *γ* across those 20 cells, each axis normalised to the extent of the morphospace. The degree and connection-distance comparison in Figure 4B and C instead uses the 20 best-fitting networks of each measure. Their distributions were compared against those of the 100 individual empirical connectomes. These two tests indicate whether a measure’s notion of similarity recovers the topological and spatial organisation of real brain networks. A small but important note: The group consensus that every measure is fitted against is not itself typical in this respect, compared to the individual empirical connectomes: at the same number of edges (495), the consensus carries a mean connection length of 48.7 mm and 9.7% of its connections beyond 90 mm, against 42.1 ± 1.1 mm and 6.6 ± 1.2% across the individuals. The two are built by different routes: each individual network keeps its 10% strongest connections, while the consensus is assembled to approximate the group’s edge-length distribution at the same density. The plausibility filter in Figure 3E instead uses the 100 best-fitting parameter combinations of each measure, and the reference-noise and parameter-recovery analyses use the single best-fitting parameter combination, i.e. the argmin of the replicate-averaged landscape.

### 4.5 Hub Topography

Generated and empirical networks share the Schaefer-100 parcellation and are therefore node-aligned, which allows for an additional topographic rather than solely distributional comparison. For every (*η, γ*) combination we averaged the nodal degree vector over its 10 replicates and correlated it, across the 100 nodes, with the nodal degree vector of the empirical consensus. The same correlation was computed for each of the 100 individual empirical connectomes against the consensus, giving the empirical reference value. In addition, each degree vector was correlated with nodal spatial centrality (the negated mean Euclidean distance from a node to all other nodes), which tests whether hub placement is explained by geometry alone. The full protocol and the results can be found in Section S6.

### 4.6 Runtime Analysis

We ran each distance measure evaluation on 25,000 networks (i.e. comparing each one of the 25,000 generated networks against the empirical consensus network), and timed the evaluation time for each of the network comparisons. Runtime is a property of the machine rather than of the networks, so all timings quoted in this work come from one reference measurement of these 25,000 comparisons, taken in a single session on one machine; they are not re-measured alongside later re-scorings of the morphospace.

### 4.7 Synthetic Parameter Recovery

#### Parameter ranges used so far

To assess the parameter recovery accuracy with known ground truth, we drew 100 parameter combinations uniformly from the whole morphospace (*η ∈* [*−*8.0, 3.0], *γ ∈* [*−*0.1, 1.0]) as ground-truth targets and generated one test network at each of them using the same GNM framework (matching index rule, MST seed). Separately, a 10 × 10 recovery grid was constructed by generating 30 networks at each of the 100 grid points (*η ∈* [*−*8.0, 3.0], *γ ∈* [*−*0.1, 1.0], linearly spaced) and turning them into a per-point consensus, following the same methodology as for the creation of the empirical consensus network. Each test network was compared against all 100 grid-point consensuses; the grid point with the minimum distance was taken as the predicted parameter combination. The recovery error was quantified as grid steps: the Euclidean distance in grid-index space between the true nearest grid cell and the predicted cell, averaged across all 100 test networks.

#### Parameter recovery within the plausible range for human connectomes

The experiment above was repeated inside the region of parameter space that the measures actually point at for human connectomes. The window was defined by pooling the locations of the 100 best-fitting networks of each of the eight selected measures against the empirical consensus (800 points in total) and taking the interquartile range of those locations, which gives *η ∈* [*−*3.96*, −*1.49] and *γ ∈* [0.08, 0.30]. It was re-derived on the corrected landscapes described in Section 4.2, and the whole experiment (i.e. ground-truth networks, recovery grid, and scoring) was regenerated inside it. From this window we drew 100 ground-truth combinations uniformly and generated one network per combination. Recovery used a dedicated 10 × 10 grid spanning *η ∈* [*−*4.31*, −*1.14] and *γ ∈* [0.05, 0.34] (which is deliberately slightly wider than the sampling window, so that recovery is not truncated at its edges) with each grid point represented by a consensus of 20 networks. Recovery was scored both in grid steps, as above, and as the Pearson correlation between true and recovered *η*. The full results are in Section S8.

#### Chance level

A measure that returns a uniformly random cell of the 10 × 10 grid has an expected grid-step error given by the mean Euclidean index distance between two independently drawn cells. On the wide grid, where the ground-truth combinations are drawn uniformly from the whole morphospace, this expectation equals 4.96 grid steps (3.16 along a single axis); inside the plausible window it equals 4.53 grid steps (2.90 along a single axis). These are the chance levels drawn as reference lines in Figure 5A and B.

### 4.8 Real Versus Synthetic Connectomes

To test whether a measure ranks real connectomes ahead of synthetic ones, we aimed to compare for each measure as how differently it evaluates both real connectomes and synthetic connectomes in comparison to the empirical consensus connectome. For this, each of the 100 individual empirical connectomes was scored against a consensus rebuilt from the other 99 subjects, so that no subject contributes to its own reference. These leave-one-out consensuses were built by the same pipeline as the main consensus (`struct_consensus` followed by binarization at 10% density), so the only difference between a leave-one-out reference and the reference used elsewhere in this work is the excluded subject. The 25,000 GNM networks were scored against the full empirical consensus connectome (generated from all 100 subjects). Discrimination was quantified as the area under the ROC curve (AUC), i.e. the probability that a randomly chosen real subject is closer to the consensus than a randomly chosen synthetic network. This analysis was computed twice: once against all 25,000 GNM networks, and once against only each measure’s 500 best-fitting networks.

### 4.9 Uncertainty-Measurements

Quantification of the uncertainty in individual measures was performed with the coefficient of variation (CV) and the intrinsic signal to noise ratio (iSNR):

Coefficient of Variation (CV)

For each individual landscape pixel (*η, γ*) with mean *µ_η,γ_* and std *σ_η,γ_* across the 10 replicates,

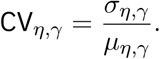

Then, the reported CV is the mean over all *η*-*γ*-combinations (i.e. all pixels *N_p_* = 2, 500):

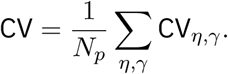

Intrinsic SNR (iSNR)

This compares the structure a landscape carries across parameter combinations to the replicate noise at each pixel. Let *M* be the mean landscape, i.e. the 50 × 50 matrix of the per-pixel means *µ_η,γ_*. The signal is the variance of *M* , we call it 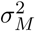. To calculate the noise, we first look at the variance across the 10 replicates, 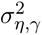. The noise is the mean 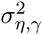 over all pixels, so 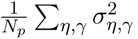. Then, the iSNR is defined as:

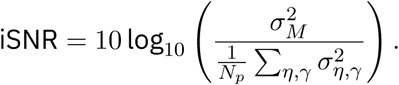

### 4.10 Degeneration Analysis

For the degeneration analysis, we disrupted the consensus network by progressively rewiring its edges and measured how each measure responded. We generated rewired networks from the original consensus as follows: at step *k ∈ {*0, 1*, . . . , K}* with *K* = 100, a rewired network was generated from the original consensus by attempting *r_k_* = *[M · k*/*K♩* double-edge swaps (NetworkX’s `double_edge_swap` function), where *M* = 495 is the number of edges. A double-edge swap exchanges the endpoints of two edges, so it preserves every node’s degree while changing which pairs of nodes are connected. Each network is generated fresh from the original consensus rather than cumulatively from the previous step, so *k* is the number of attempted swaps, i.e., *k* = 0 returns the unperturbed consensus and *k* = *K* attempts one swap per edge.

The number of attempted swaps is not the number of edges that change. `double_edge_swap` rejects any swap that would create a self-loop or duplicate an existing edge, and a later swap can move an edge that an earlier one already moved. To illustrate, attempting one swap per edge of the consensus changes only 78% of them, and a response curve plotted against attempted rather than effective change looks considerably more non-linear than it is. Thus, we report all results against the *effective degeneration*, the Hamming distance 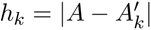 between the original adjacency matrix *A* and the rewired adjacency matrix 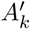 , which counts the edges that actually differ. *h_k_* thus grows with *k* in expectation, but saturates below *M* , as the degree-preserving randomisation keeps some of the original edges by chance.

To account for stochasticity in the rewiring, we generated 200 independent rewiring trajectories, each starting from the original consensus with a different random seed, and evaluated every distance measure on the same 200 trajectories, so that differences between measures are not confounded with differences in the networks they were computed on.

To assess how faithfully each distance measure tracks effective structural disruption, we computed the mean absolute error (MAE) between each measure and Hamming distance, which served as a reference measure of true edge-level change. Prior to computing the MAE, both the measure values and the Hamming distances were min-max normalised to [0, 1] within each trajectory. The MAE for each measure was then computed as:

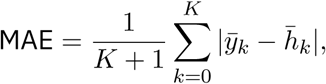

where *ȳ_k_* and *h̄_k_* are the measure value and the effective degeneration at level *k*, each min-max normalised per trajectory and then averaged over the 200 trajectories. These are exactly the curve and the diagonal drawn in Figure 4D, so the MAE in Figure 4F is the area between them. A lower MAE indicates that the measure’s response to rewiring closely mirrors the actual number of changed edges, while a higher MAE indicates a non-linear or otherwise distorted sensitivity profile.

Stability under rewiring was quantified with a second coefficient of variation that differs slightly from the landscape CV defined in Section 4.9: at each perturbation level *k* we took the standard deviation of the measure’s value across the 200 trajectories divided by its mean at that level, and report the mean of that ratio over all levels. Measures defined as similarities were converted with the min-max-and-invert convention of Section 4.4 before this ratio was formed. The values reported in the Results (Frobenius 0.0130, DeltaCon 0.0161, and so on) are this rewiring CV.

### 4.11 Edge Importance

For each of the 495 edges of the consensus in turn, the edge was removed and the distance between the resulting network and the unperturbed consensus was computed. The resulting increase is that edge’s importance under that measure. Figure 4E shows the top 20% of edges by importance for each measure, drawn at the anatomical coordinates of the Schaefer-100 parcellation.

### 4.12 Robustness of the Recovered Parameters to Reference Noise

The degeneration analysis above perturbs the networks being compared, while the analysis described in this Section perturbs the reference and evaluates the drift in the recovered parameter combination. The consensus was rewired by *s ∈ {*1, 2, 5, 10, 20, 50, 100, 200, 495*}* connected degree-preserving double-edge swaps, with 10 to 20 repeats per level depending on the cost of the measure. At each level the best-fitting (*η, γ*) combination was re-recovered on the 50 × 50 landscape and its drift from the unperturbed best-fitting combination was measured in grid steps. A measure’s noise tolerance *N^∗^* is the largest perturbation level at which its median drift is still within one grid step, reported as the effective degeneration of that level (the fraction of the 495 consensus edges that actually differ), as in the degeneration analysis above. The full results are in Section S9.

### 4.13 Summary comparison of the selected measures

The radar plots in Figure 1D summarize each measure on the five axes the Results are structured around. Each axis is represented by the single read-out that reduces most cleanly to one number per measure: *agreement* is the mean Pearson correlation of a measure’s landscape with the landscapes of the other seven selected measures (from the matrix in Figure 3B); *biological plausibility* is the total variation of the locations of the measure’s 20 best-fitting parameter combinations (Figure 4A); *computational efficiency* is the mean runtime per comparison (Figure 3D); *sensitivity and robustness* is the mean of three ranks, one per perturbation read-out: the MAE between the rewiring response and the diagonal (Figure 4D and F), the coefficient of variation across the 200 rewiring trajectories (Figure 4G), and the noise tolerance *N^∗^* of the recovered parameters (Section S9); the first covers sensitivity, the other two robustness; and *accuracy* is the mean grid-step error inside the plausible parameter window (Figure 5B). Runtime and grid-step error are lower-is-better quantities and were inverted, so that on every spoke a larger radius denotes the more desirable value.

Then, we assigned scores by ranking and comparing the measures: on four of the five axes the measures are ordered by their underlying value and given their *rank as the score* (8 as most desireable value down to 1 as the least; if measures tie their ranks get averaged). The accuracy axis is scored against its chance level. Also, because a recovery error at or beyond chance carries no information about the parameters, these measures get scored as 0. All measures performing better than chance are spread linearly between 0 and 8. Three of the eight measures (KS-based energy, spectral (non-backtracking), and spectral (adjacency)) score 0 on this axis.

Two notes about these plots follow from the scoring by each measure’s rank: First, while the scoring preserves the ordering of the measures on each axis, the four rank-scored axes do not preserve the size of the differences (which on some axes span orders of magnitude). Second, the scores are relative to this set of eight measures, so adding or removing a measure changes them.

### 4.14 Software

All analyses were implemented in Python. Libraries used throughout are `NetworkX` ^48^, `netrd`^49^, `netneurotools`^44^, `generativenetworkmodels`^36^, `scikit-learn`^50^, `SciPy`^51^, and `NumPy` ^52^. The code and the derived data are available as stated in the Data and Code Availability statement.

## DATA AND CODE AVAILABILITY

The structural connectome data underlying this study come from the Human Connectome Project S900 release^40^ and were processed as part of^39^; they are available from the Human Connectome Project subject to its data use terms. The second dataset used in the Supplementary Material is the Human Connectome Project Young Adult release^53^, accessed through DSI Studio’s Fiber Data Hub^54^. All analysis code, together with the generated networks and the 16 distance landscapes that underlie every figure, is available at https://github.com/ad045/14_4D_benchmarking.

## ACKNOWLEDGEMENTS

A large language model (Anthropic Claude, via Claude Code) was used during this project to assist with software development and for quality assurance. Additionally, it provided language and editorial support during the manuscript preparation. All model-assisted output was reviewed, tested, and edited by the authors, who are solely responsible for the final content of the manuscript.

## AUTHOR CONTRIBUTIONS

Conceptualization: A.D., K.F., D.E.A. Methodology: A.D., K.F., A.I.L., F.P. Software: A.D. Formal analysis: A.D., K.F. Investigation: A.D., K.F. Data curation: A.M., A.I.L., K.F., A.D. Visualization: A.D., K.F. Writing - original draft: A.D., K.F. Writing - review and editing: all authors. Supervision: K.F., D.E.A.

