## Supplementary material for "How Similar Are Two Brains? A Comprehensive Benchmark of Brain Network Similarity Measures"

Supplementary figures and tables are numbered S1, S2, etc; unprefixed references to 'Fig.', 'Table', 'Section', or 'Methods' point to the main article.

#### S1 Network Distance Measures

This section provides an overview of all 16 measures that were evaluated. They are listed with their defining formula and source in Table S1; afterwards, the less common ones are explained in more detail. We use the term *distance measure* throughout, noting that several of the 16 do not qualify as metrics in the mathematical sense, as they lack the formal properties a metric must fulfil[1]. All measures were computed on binarized undirected graphs, even where they allow for weighted or directed ones. As node correspondence is known for a given cortical parcellation, we include measures that require it, such as DeltaCon.

Throughout, a graph is written $G = (V, E)$ with node set $V$ of size $n = |V|$ and edge set $E$, and $A$ denotes its $n \times n$ adjacency matrix with entries $a_{ij} = 1$ if $\{i, j\} \in E$ and $a_{ij} = 0$ otherwise. The second graph of a compared pair, its edge set, and its adjacency matrix are marked with a prime ($G'$, $E'$, $A'$).

For visualization and comparison, all measures were min-max-normalized to $[0, 1]$. Some of the 16 are defined as distances and others as similarity scores; similarity scores were first normalized ($s_{norm}$) and then turned into normalized distances by

$$d_{norm} = 1 - s_{norm}.$$

Table S1 gives each measure as it is typically defined, i.e. as a similarity or as a distance.

| Measure Name | Type | Key Formula | Citation |
|---|---|---|---|
| Frobenius | Matrix | $\|A - A'\|_F$; on binarized undirected graphs $= \sqrt{2\sum_{i<j} \mathbf{1}[a_{ij} \neq a'_{ij}]}$, the square root of twice the number of differing edges | Linear algebra |
| Hamming | Edge | $\sum_{i<j} \mathbf{1}[A_{ij} \neq A'_{ij}]$ | 2 |
| Jaccard | Edge | $\|E \cap E'\| / \|E \cup E'\|$ | Set theory |
| F1 | Edge | $2PR / (P + R)$, with precision $P$ and recall $R$ | Statistics |
| Network Mutual Information (NMI) | Information | $2I(G; G') / (H_b(G) + H_b(G'))$, with mutual information $I$ and binary entropy $H_b$ of the vectorized adjacency matrix of $G$ | 3 |
| Degree-Corrected NMI | Information | $2I_{\text{DC}} / \left(n^{-1} \sum_{i=1}^{n} H_b(p_i)\right)$; like NMI, but conditioned on the degree sequence, so the normalization runs over the per-node connection probabilities $p_i$ instead of the graph-wide edge density | 3 |
| Spectral (adjacency) | Spectral | $\sqrt{\sum_{i=1}^{n} (\lambda_i^A - \lambda_i^{A'})^2}$, with $\lambda_i^A$ the $i$-th largest eigenvalue of $A$ | 4 |
| Spectral (norm. Laplacian) | Spectral | Same as spectral distance (adjacency), just on $L_{\text{norm}} = D^{-1/2} L D^{-1/2}$ instead of on $A$ | 5 |
| Spectral (non-backtracking) | Spectral | Same as spectral distance (adjacency), just on non-backtracking matrix $B$ instead of on $A$ | 6 |
| Portrait divergence | Feature | JSD$(P \parallel P')$, with the Jensen-Shannon divergence JSD, a symmetrized and bounded version of the Kullback-Leibler divergence, and $P$ and $P'$ being the portraits of $A$ and $A'$, respectively | 7 |

| Measure Name | Type | Key Formula | Citation |
|---|---|---|---|
| KS-based energy | Feature | $\max(\mathsf{KS}(k), \mathsf{KS}(c), \mathsf{KS}(b), \mathsf{KS}(\ell))$, over the distributions of degree $k$, clustering coefficient $c$, betweenness centrality $b$, and edge length $\ell$ | 8 |
| NetSimile | Feature | $\sum_k \lvert f_k - f'_k \rvert \,/\, (\lvert f_k \rvert + \lvert f'_k \rvert)$, with $f_k$ and $f'_k$ being the feature vectors of $A$ and $A'$ | 9 |
| DeltaCon | Kernel | $1 \,/\, (1 + d_{\mathsf{M}}(S, S'))$, with $d_{\mathsf{M}}$ the Matusita distance between the node-influence matrices $S$ and $S'$ | 10 |
| Communicability JSD | Communi-cability | JSD on normalized Communicability, with the Jensen-Shannon divergence JSD, a symmetrized and bounded version of the Kullback-Leibler divergence | 11 |
| Communicability Correlation Distance | Communi-cability | Pearson correlation between Communicability | Custom |
| Resistance | Resistance | $\lVert R - R' \rVert_2$, where $R$ and $R'$ are the resistance matrices of $A$ and $A'$ respectively | 12 |

Table S1: **Overview of the 16 network distance measures evaluated.** For each measure its category, the formula defining it, and its source are given. $A$ and $A'$ are the adjacency matrices of the two compared graphs.

The following will provide more information on less known distances and similarities. For the eight measures selected in the main text, the description is followed by a figure that displays the intermediate steps of that measure on a single pair of graphs. The figure for DeltaCon is located in the main text. The pair of example networks that are compared in the displays is the same in all eight figures: $G_1$ is a ring lattice of 24 nodes in which every node is joined to its three nearest neighbours on either side (72 edges); $G_2$ is that same lattice with 15% of its edges rewired to random endpoints, drawn in orange. Both graphs have the same number of nodes and edges, so the two differ only in where the connections sit; thus, this setup represents the situation a distance measure faces when it compares a generated network to an empirical connectome at matched density.

Every figure follows the same layout: The upper row shows $G_1$ and the lower row $G_2$, each drawn as a graph and as its adjacency matrix $A$, followed by the representation the measure builds from $A$ (e.g., a node-influence matrix, an eigenvalue spectrum, a communicability matrix, a portrait, a feature table, or a set of property distributions). Arrows lead from the two rows into the panel in which the two representations are combined, and the resulting distance is printed underneath it. The numbers shown are the values for this specific pair of graphs and are not comparable across measures, since the measures have different ranges; within the benchmark all distances are min-max normalized as described above.

#### Matrix-based measures

The Frobenius distance is defined in more detail below. Hamming, Jaccard, and F1 distance are defined in Table S1.

**Frobenius distance** The Euclidean norm of the element-wise difference of the two adjacency matrices,

$$d_{Frob}(G, G') = \|A - A'\|_F = \sqrt{\sum_{i=1}^{n}\sum_{j=1}^{n}(a_{ij} - a'_{ij})^2}.$$

On binarized undirected graphs every entry of the difference is either 0 or ±1, so the sum counts the entries on which the two matrices disagree and $d_{Frob}$ is the square root of twice the number of differing edges.

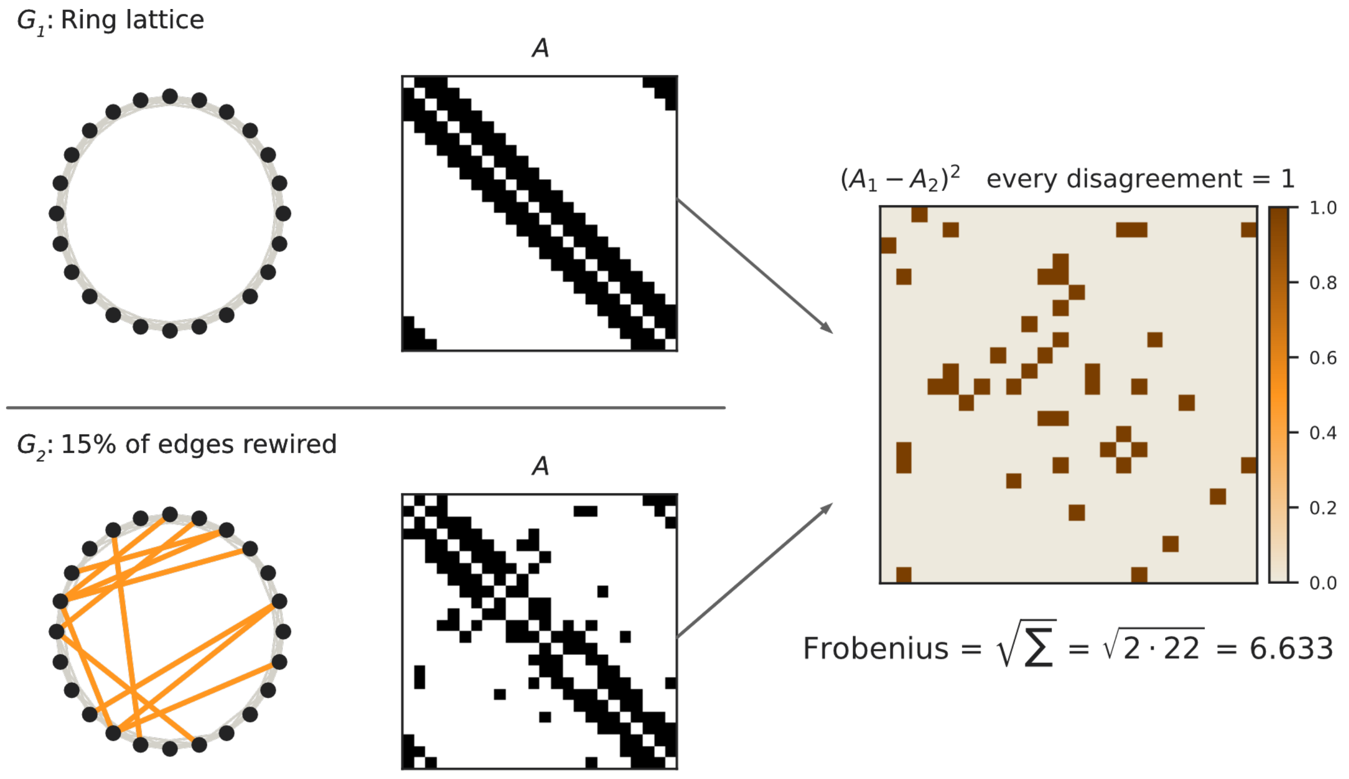


Figure S1: **Frobenius distance.** Left: $G_1$, $G_2$, and their adjacency matrices. Right: the element-wise squared difference $(A_1 - A_2)^2$, in which every entry on which the two matrices disagree takes the value 1. The distance printed below is the square root of the sum over all entries.

#### 50 Information-theoretic measures

**Network Mutual Information (NMI)** Based on Shannon's mutual information[3]. Each graph $G$ is represented by the vectorized upper triangle of its adjacency matrix, a binary vector treated as a random variable, so the shared information is measured at the level of individual edges. Then, $MI(G;G')$ is the mutual information of the two edge vectors and $H_b(G)$ the binary entropy of the edge density $p$ of $G$:

$$NMI(G,G') = \frac{2 \cdot MI(G;G')}{H_b(G) + H_b(G')},$$

with

$$H_b(p) = -p \log_2(p) - (1-p) \log_2(1-p).$$

51 The implementation stems partly from `network-MI`[13].

**Degree-corrected NMI (DC-NMI)** NMI variant that accounts for degree sequence preservation[3], and thus takes the degree heterogenity of graphs into account:

$$NMI_{DC}(G,G') = \frac{2 \cdot MI_{DC}(G;G')}{\frac{1}{n}\sum_{i=1}^{n}[H_b(p(i)) + H_b(p'(i))]}.$$

52 Again, the implementation is based on[13].

#### 53 Spectral measures

**Spectral distance (Adjacency)** Euclidean distance between adjacency matrix eigenvalue spectra with the eigenvalues $\lambda_i^A$ and $\lambda_i^{A'}$:[4]

$$d_{ASD}(G,G') = \sqrt{\sum_{i=1}^{n}(\lambda_i^A - \lambda_i^{A'})^2}.$$

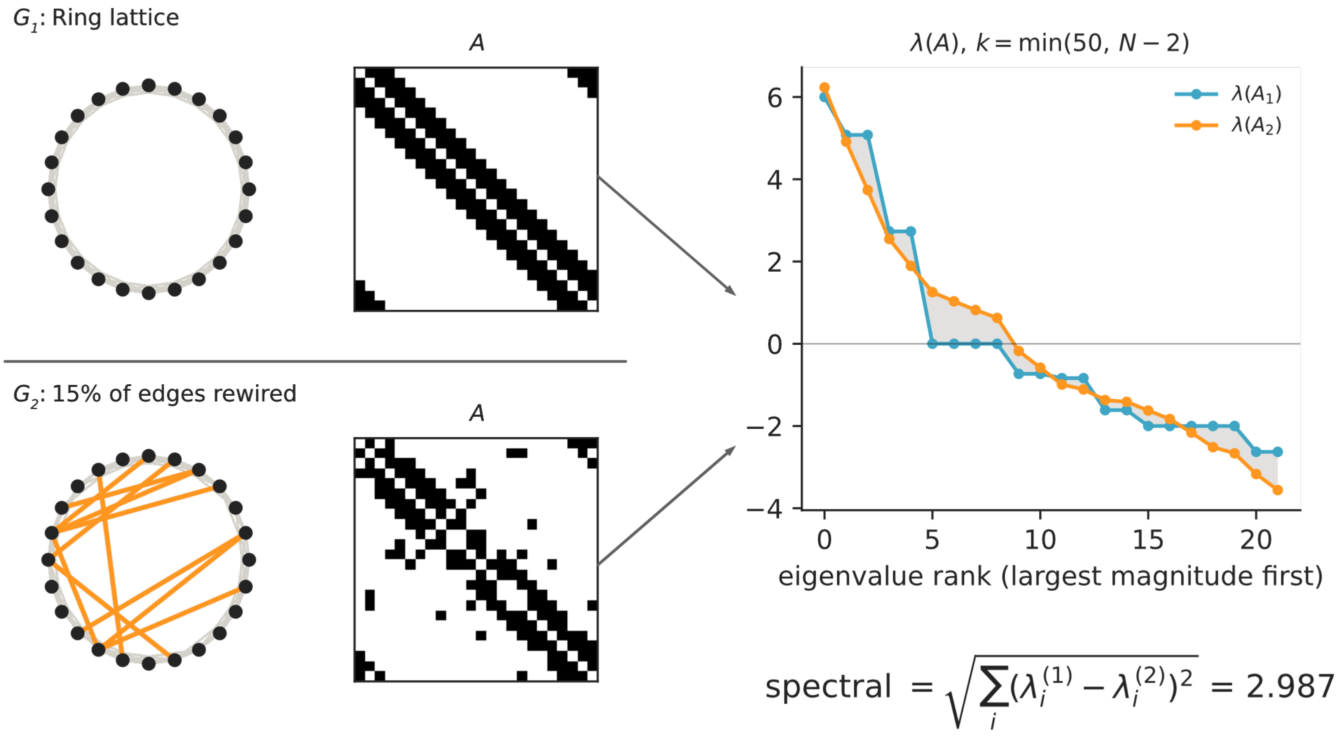


Figure S2: **Spectral distance (adjacency).** Left: $G_1$, $G_2$, and their adjacency matrices. Right: the leading $k = \min(50, N - 2)$ eigenvalues of $A_1$ and $A_2$, ordered by decreasing magnitude; the shaded area marks the gap between the two spectra. The value printed below is the Euclidean distance between the two eigenvalue sequences.

**Spectral distance (Normalized Laplacian)** Distance between normalized Laplacian eigenvalues; the normalized Laplacian is defined as $L = D^{-1/2}LD^{-1/2}$.

**Spectral distance (Non-Backtracking)** Distance based on non-backtracking matrix (shape: $m \times m$ for undirected graphs, where $m$ is the number of edges) eigenvalues[6]. This work uses the implementation of `netrd`[14].

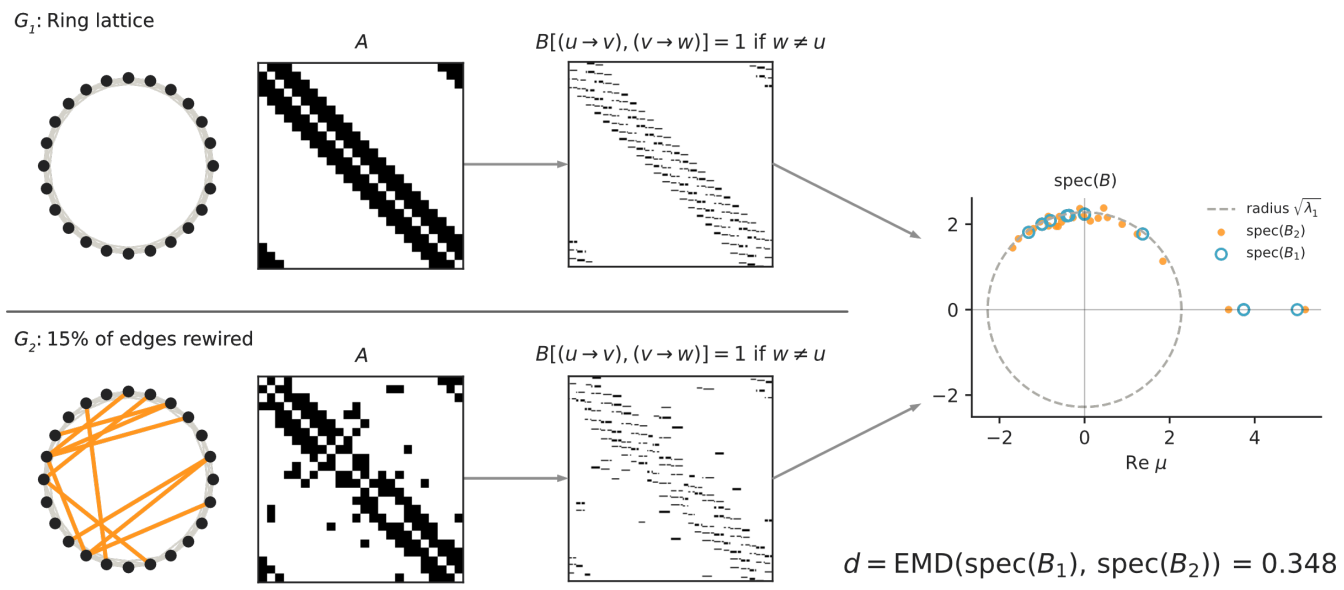


Figure S3: **Spectral distance (non-backtracking).** Per row: the graph, its adjacency matrix $A$, and the non-backtracking matrix $B$, indexed by directed edges, with $B[(u \to v), (v \to w)] = 1$ whenever $w \neq u$. Right: the eigenvalues of $B_1$ (open circles) and $B_2$ (filled dots) in the complex plane; the dashed circle has radius $\sqrt{\lambda_1}$. The value printed below is the earth mover's distance between the two spectra.

### Graph kernel and feature-based measures

**Portrait Divergence** Jensen-Shannon divergence between network portrait matrices[7].

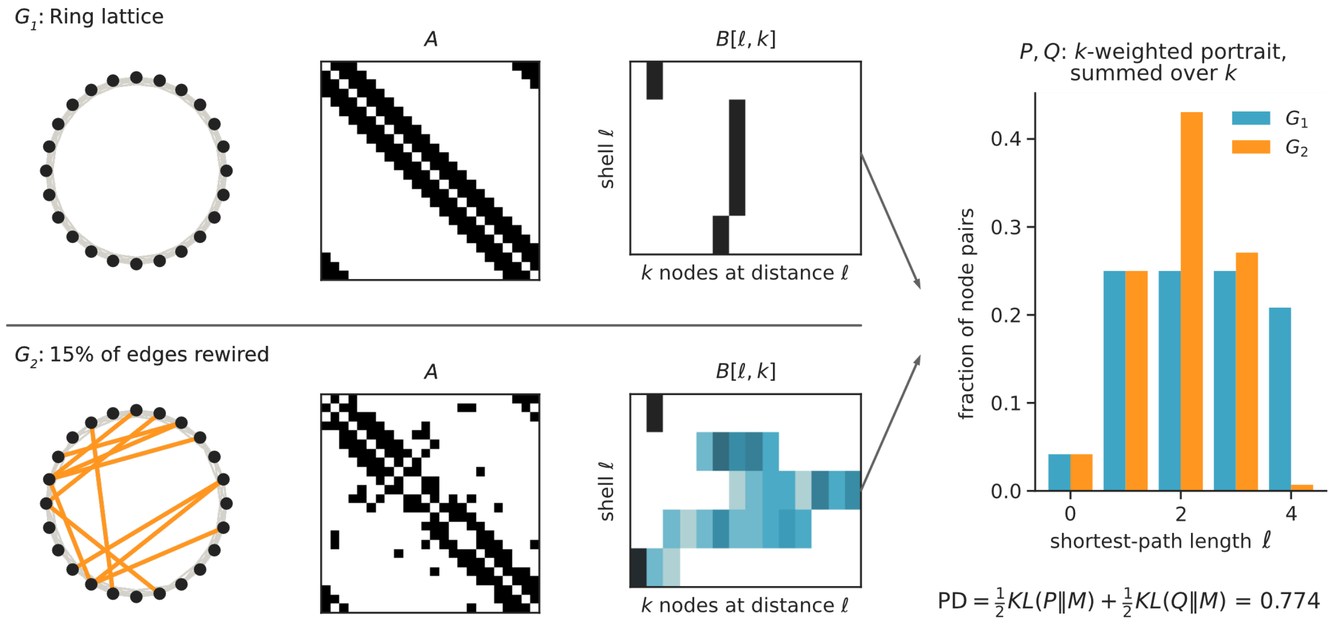


Figure S4: **Portrait divergence.** Per row: the graph, its adjacency matrix $A$, and the network portrait $B[\ell, k]$, the number of nodes having exactly $k$ nodes at shortest-path distance $\ell$. Right: the two portraits reduced to the distributions $P$ and $Q$ over shortest-path lengths by $k$-weighted summation over $k$. The value printed below is the Jensen-Shannon divergence between $P$ and $Q$, with $M$ their mean.

**NetSimile** The Canberra distance between the feature vectors of the two graphs[9]; for the definition see Figure S5. The feature vectors contain the mean, median, standard deviation, skewness, and kurtosis of the following descriptors, based on the node itself (its degree; its clustering coefficient: i.e. how many triangles this node is part of), on its neighbors (their average degree; their average clustering coefficient), and on its egonet (its number of edges; its number of outgoing edges; its number of neighbors); resulting in a vector with $7 \cdot 5$ entries. In this work, the implementation of `netrd`[14] was used.

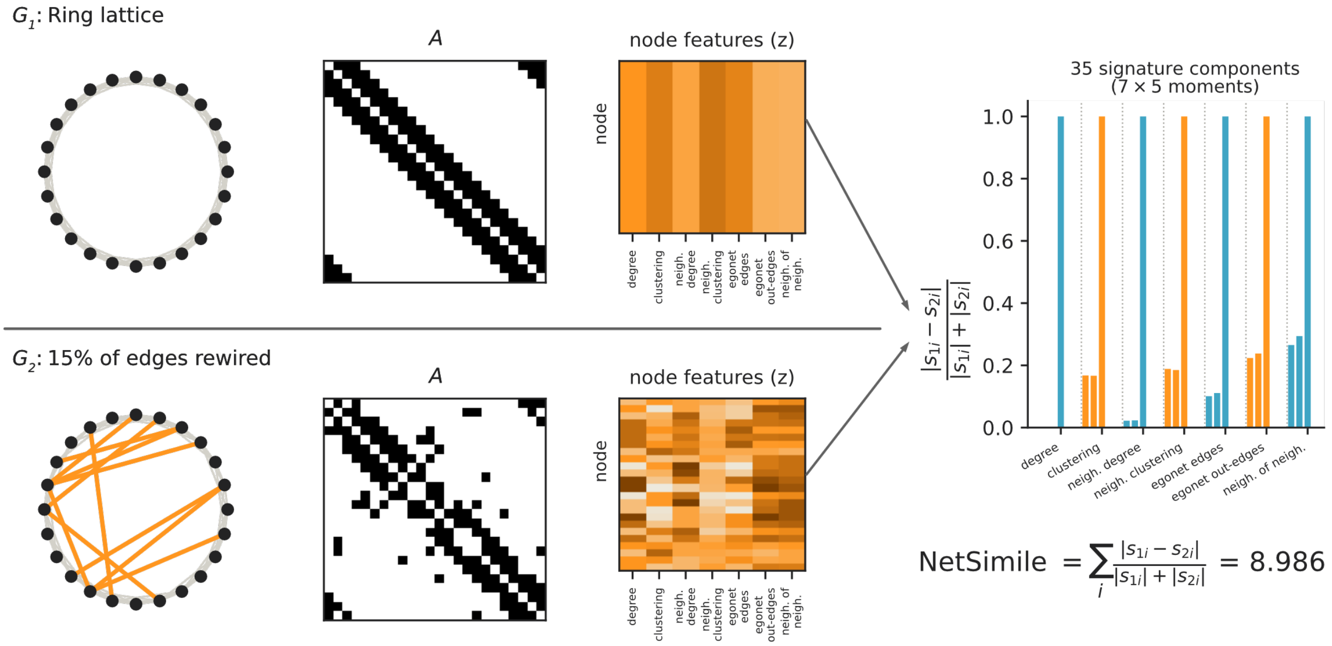


Figure S5: **NetSimile.** Per row: the graph, its adjacency matrix $A$, and the table of the seven node features (z-scored, one row per node). Right: the per-component Canberra terms $|s_{1i} - s_{2i}|/(|s_{1i}| + |s_{2i}|)$ of the 35-entry signature, which holds five distributional moments per feature; bar colour alternates between features and dotted lines separate them. The value printed below is the sum over all 35 components.

**DeltaCon** This similarity measure is based on fast belief propagation comparing node affinities[10]. For this, similarity matrices $S_{ij}$ - modelling the influence flow through the network - are calculated: Direct connections (i.e. 1-hop neighbors) that show the strongest influence and indirect paths (2-hop, 3-hop, etc) that show weaker influence that decays with distance. The measure looks at all possible paths, capturing that if nodes are connected by more paths that this then leads to stronger influence between them. In other words, the calculation of $S$ is based on 'fast belief propagation':

$$S = [I + \epsilon^2 D - A]^{-1},$$

where $I$ is the identity matrix, $D$ is the diagnoal degree matrix, and $\epsilon$ is a small constant that describes neighbour influence.

The similarity between two graphs is then calculated using the Matusita distance (the root Euclidian distance - a measure that boosts small differences and thus increasing similarity of small changes) between the similarity matrices $S$,

$$d_{Matusita} = \sqrt{\sum_{i=1}^{n}\sum_{j=1}^{n}(\sqrt{s_{1,ij}} - \sqrt{s_{2,ij}})^2},$$

with then leads to the computation of a final distance:

$$DeltaCon(G, G') = \frac{1}{1 + d_{Matusita}}.$$

The value of looking at node similarities can be illustrated by a small example: Take a barbell graph (i.e. a graph that has two cliques that are connected by a single bring edge). If now one random edge within a clique is removed, then information can still flow through many alternative paths. If, however, the bridge is removed, and thus the graph is disconnected, information ceases to flow. DeltaCon recognizes the bridge as far more important because it dramatically changes how influence propagates - which is not the case of simple edge-counting measures.

Scalability of this algorithm is achieved by approximation, that partitions nodes into $g$ groups and that then computes group affinities rather than all pairwise node affinities. This reduces the complexity to $O(g \cdot m)$, where $m$ is the higher number of edges between the two graphs. The $g$ calculations are independent from each other and can be calculated in parallel. DeltaCon requires node correspondance, and captures both local and global structure. The approach has been evaluated to be helpful in comparative studies (e.g. in[1]). This work uses the implementation from `netrd`[14].

The figure visualizing the step-by-step process of calculating DeltaCon is shown in the main article (Figure 2).

**KS-based energy distance** As described in the main article, this measure combines Kolmogorov-Smirnov statistics across multiple network property distributions (degree, clustering coefficient, betweenness centrality, edge length)[8].

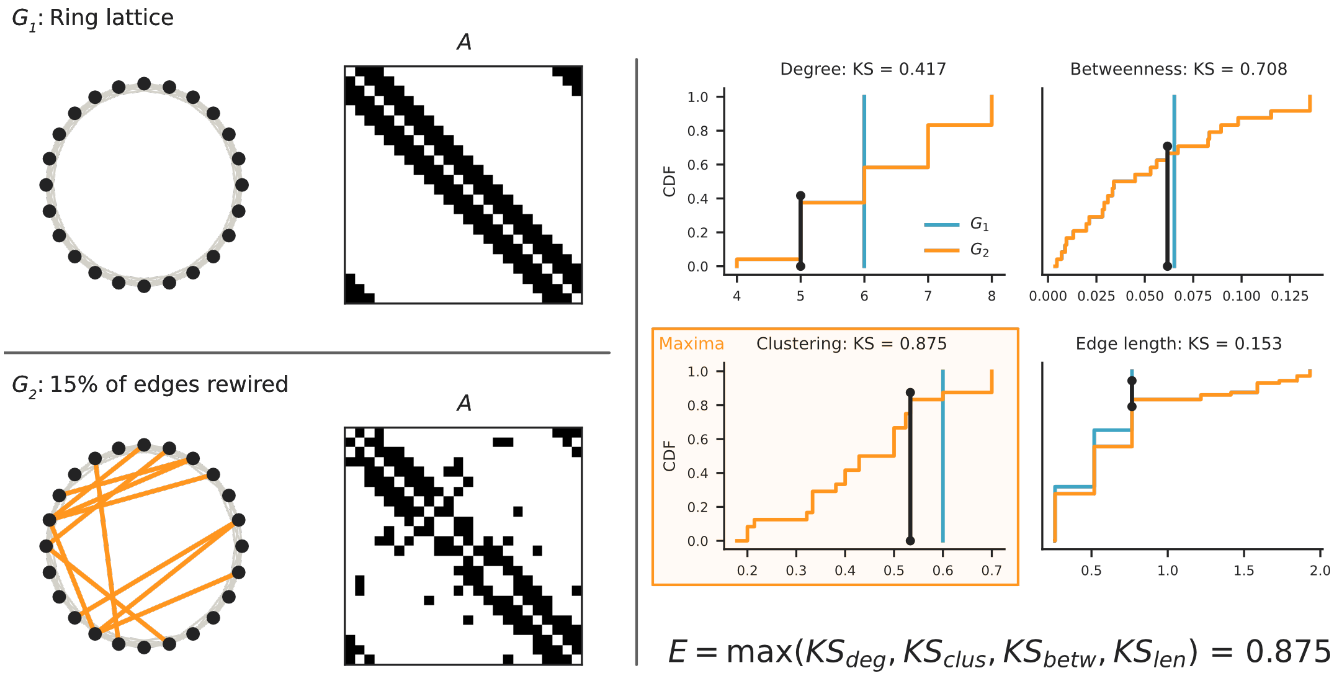


Figure S6: **KS-based energy.** Left: $G_1$, $G_2$, and their adjacency matrices. Right: the cumulative distributions of the four network properties for $G_1$ (blue) and $G_2$ (orange); the black bar in each panel marks the point of maximum vertical separation, i.e. the Kolmogorov-Smirnov statistic of that property. The panel supplying the largest of the four is framed in orange. The value printed below is that maximum.

#### 86 Communicability and resistance measures

**Communicability Correlation** This measure first maps each network to its communicability matrix $C = \exp(A)$, and then correlates the two matrices entry by entry:

$$d_{\text{ComCorr}} = \text{Pearson\_Correlation}(\text{vectorize}(e^A), \text{vectorize}(e^{A'})).$$

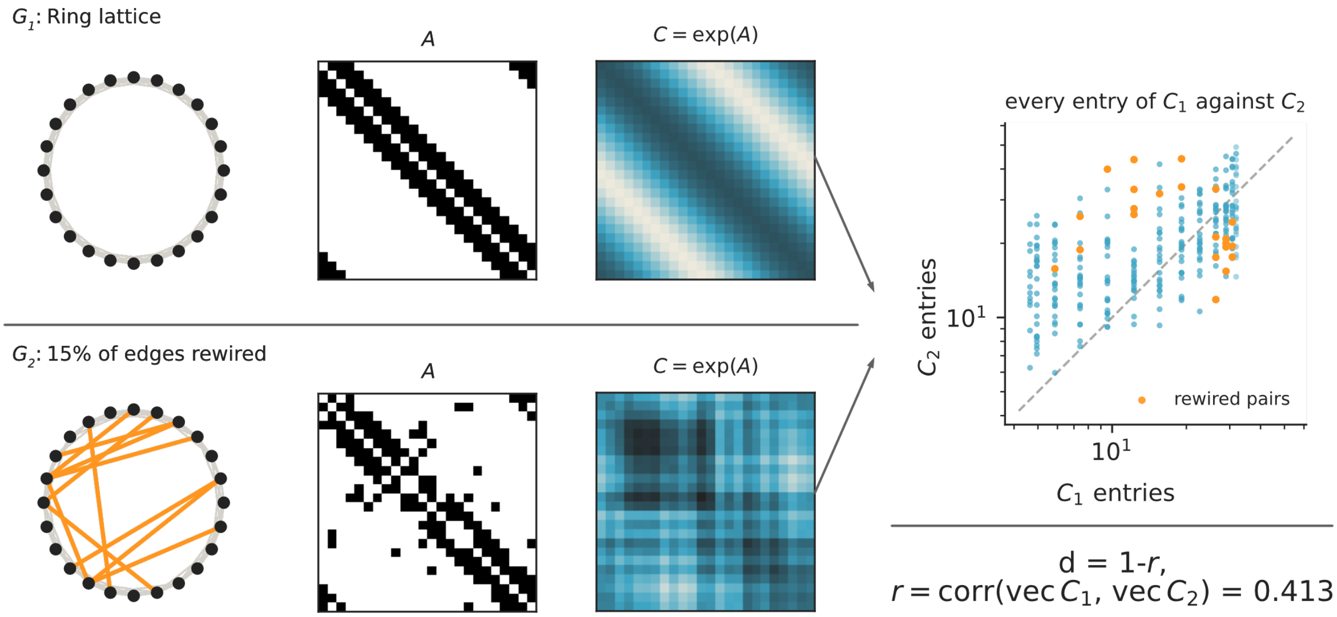


Figure S7: **Communicability correlation.** Per row: the graph, its adjacency matrix $A$, and the communicability matrix $C = \exp(A)$. Right: every entry of $C_1$ plotted against the corresponding entry of $C_2$ on logarithmic axes, with node pairs affected by the rewiring in orange and the identity line dashed. The value printed below is $1 - r$, with $r$ the Pearson correlation between the vectorized matrices.

**Communicability JSD** Jensen-Shannon divergence between communicability matrices. Normalized matrix exponentials are flattened and normalized to create probability distributions,

$$P = \frac{\text{vectorize}(e^{A'})}{|\text{vectorize}(e^{A'})|_1} \text{ and } P' = \frac{\text{vectorize}(e^{A})}{|\text{vectorize}(e^{A})|_1},$$

Then, the squared JSD with base 2 is calculated:

$$\text{JSD}(P|P') = \frac{1}{2}D_{KL}(P|M) + \frac{1}{2}D_{KL}(P'|M), \text{ where } \quad M = \frac{P+P'}{2}.$$

**Resistance distance** The resistance distance (also called the resistance perturbation distance) is calculated by comparing the effective resistance matrices of two graphs. The effective resistance matrix of a graph is defined in analogy to an electrical circuit, where each edge is modelled as resistance - the weight of it is used to define the resistance. In a binary graph, each edge could thus correspond to a resistance of 1 $\Omega$. The resistance matrix is defined as[12]:

$$R_{ij} = L^{\dagger}_{ii} + L^{\dagger}_{jj} - 2L^{\dagger}_{ij},$$

where $L^{\dagger}$ is the Moore-Penrose pseudoinverse of the Laplacian matrix $L = D - A$.

$$d_{r(p)} = \left\| R^{(1)} - R^{(2)} \right\|_p = \left( \sum_{i,j \in V} \left| R^{(1)}_{i,j} - R^{(2)}_{i,j} \right|^p \right)^{1/p},$$

87 where $p = 2$ is used in this work. This work uses the implementation of `netrd`[14].

#### S2 Landscapes: Different Datasets & Seeds

This section visualizes different KS-based energy landscapes for the 50 × 50 grid of parameter combinations, where the GNMs were generated with different seeds. This was done to show the influence of the seed choice on the resulting networks (which influence the distance to the consensus network and thus the landscape). The generated networks were compared to a consensus connectome; the consensus was generated from a dataset not used in the main manuscript: Briefly, structural connectomes from 1065 subjects from the Human Connectome Project Young Adult dataset[15] were used. These connectomes were accessed from DSI Studio's Fiber Data Hub[16] following GQI-based reconstruction[17]; the reconstruction was performed by the data providers and not by us. The connectomes were parcellated according to the Schaefer-100 atlas[18]. Connectivity was quantified using count-pass streamline counting, in which a connection is registered for every region a streamline passes through. Tractography parameters were: maximum fiber length 250 mm, minimum fiber length 30 mm, $5 \times 10^6$ streamlines, random seeding, step size 1 mm, and turning angle threshold 35°. The resulting connectivity matrices were binarized and thresholded to 10% density using `binarize_network` from `netneurotools`[19]. A consensus network was built from these networks with the same approach as described in the main manuscript.

A morphospace spanned by 25,000 GNMs (10 networks for each of the mentioned 50 × 50 parameter combinations) was generated. Then, the KS-based energy was used to determine the distance of each of the networks to the consensus connectome generated on the described second dataset. The networks were generated either with an MST seed (see Figure S8), a ring seed (see Figure S9), or no seeding (see Figure S10); again, darker colors indicate lower energy values reflecting higher similarity between the generated and consensus networks under the KS distance measure. These three runs predate the seed correction described in Section 4.2: the edges contributed by the seed were not subtracted from the number the model adds, so the MST-seeded networks carry 594 edges (a density of 12%) and the ring-seeded ones 695 (14%), against the 10% density of this dataset's consensus; only the seed-free networks sit at the matched 495 edges. This section compares seeds with one another, and each landscape is read on its own scale, so the offset does not affect the point made here - but it does mean that the absolute energy values, and the positions of the minima, are not directly comparable with those of the main analyses or between the three panels. Two notes to the seeding methods: The ring network provides a second fundamental method to connect all network nodes. For the seed-free generation, it is important to note that these networks are sometimes strongly disconnected, in some cases splitting into as many as 50 separate subgraphs.

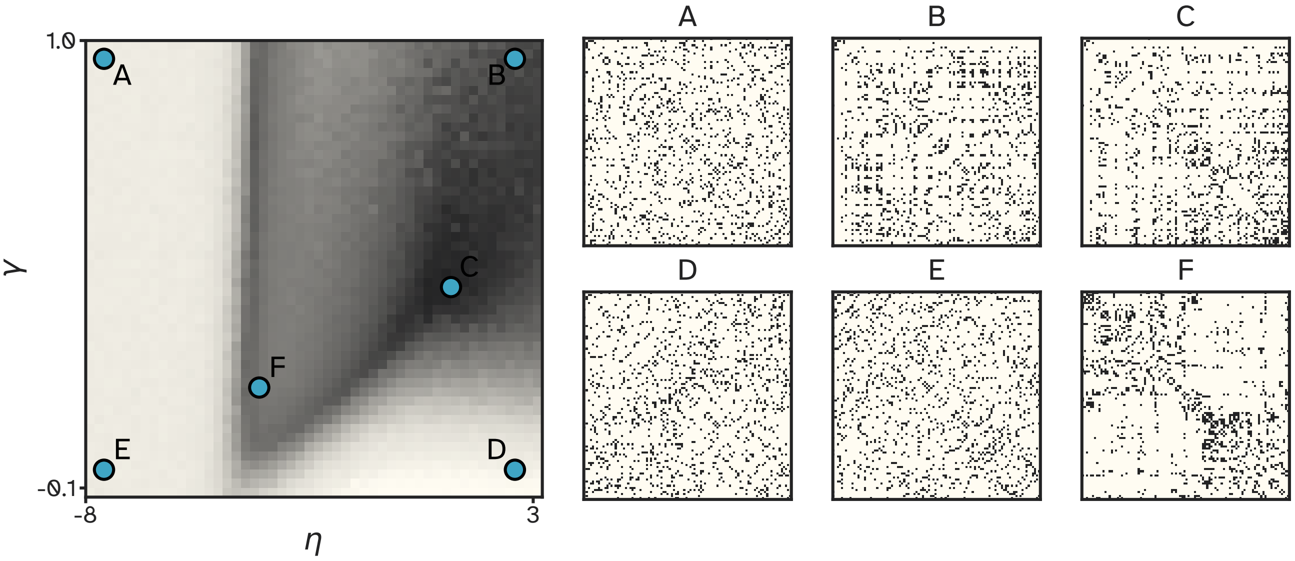


Figure S8: **Landscape on the second dataset, MST seed.** Networks were generated with an MST seed. Left: KS-based energy landscape over the 50 × 50 grid of $(\eta, \gamma)$ combinations, each cell averaged across the 10 networks generated at that combination and scored against the consensus connectome of the second dataset described in this section; darker cells indicate lower energy. Right: binary adjacency matrices of one generated network at each of the six grid points marked A-F in the landscape.

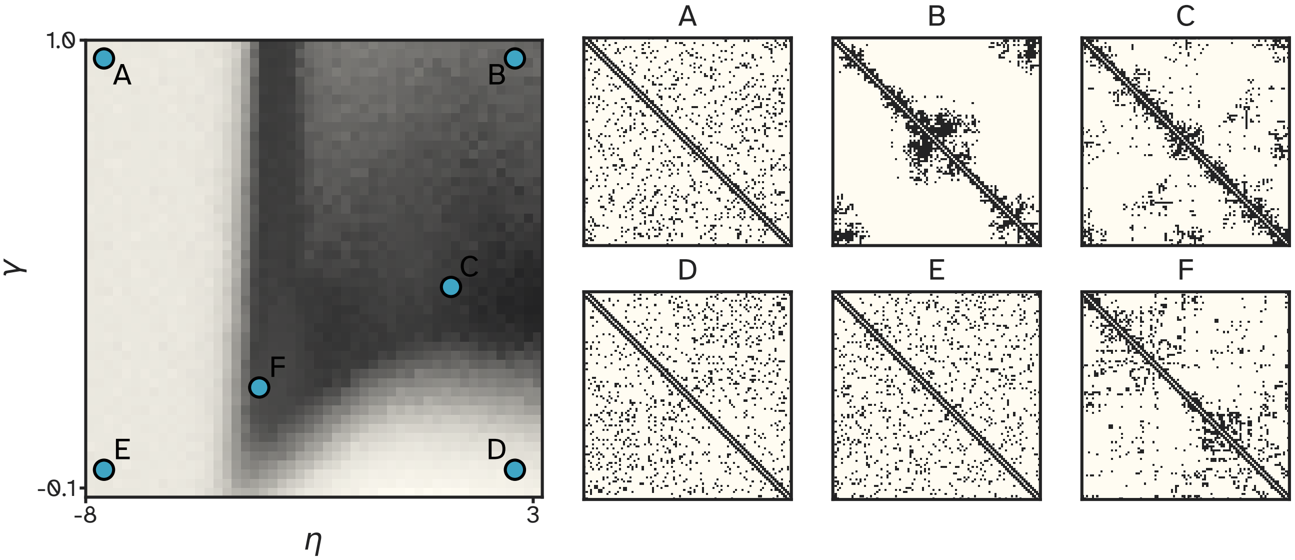


Figure S9: **Landscape on the second dataset, ring seed.** Networks were generated with a ring seed. Left: KS-based energy landscape over the 50 × 50 grid of $(\eta, \gamma)$ combinations, each cell averaged across the 10 networks generated at that combination and scored against the consensus connectome of the second dataset described in this section; darker cells indicate lower energy. Right: binary adjacency matrices of one generated network at each of the six grid points marked A-F in the landscape.

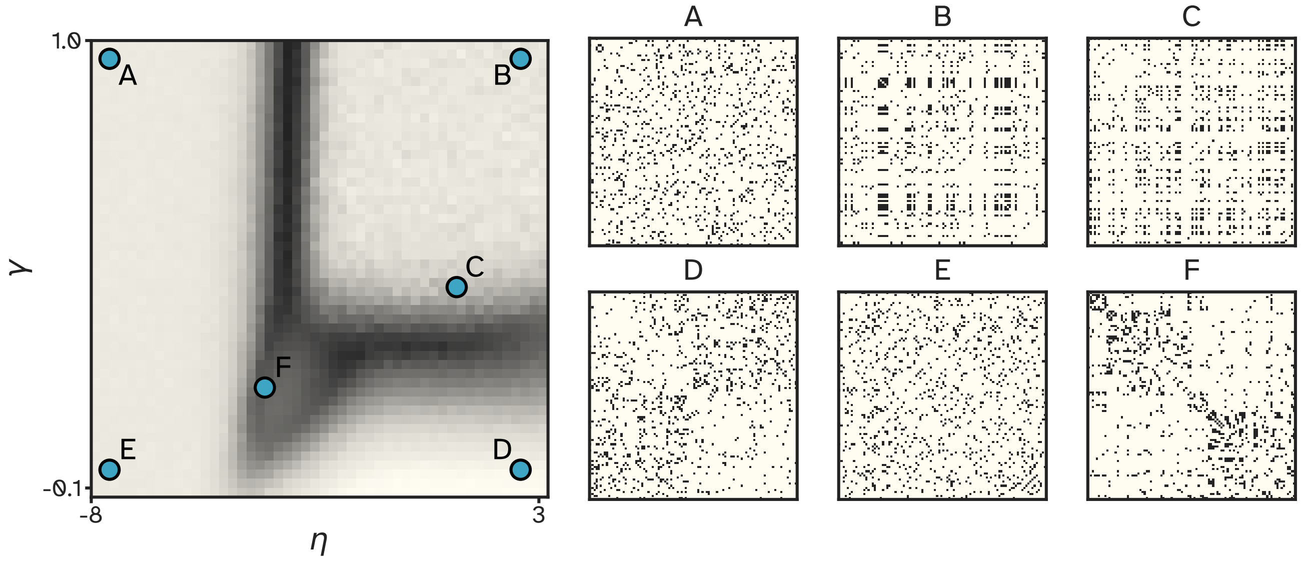


Figure S10: **Landscape on the second dataset, no seed.** Networks were generated without a seed. Left: KS-based energy landscape over the 50 × 50 grid of $(\eta, \gamma)$ combinations, each cell averaged across the 10 networks generated at that combination and scored against the consensus connectome of the second dataset described in this section; darker cells indicate lower energy. Right: binary adjacency matrices of one generated network at each of the six grid points marked A-F in the landscape.

#### S3 Components of the KS-Based Energy

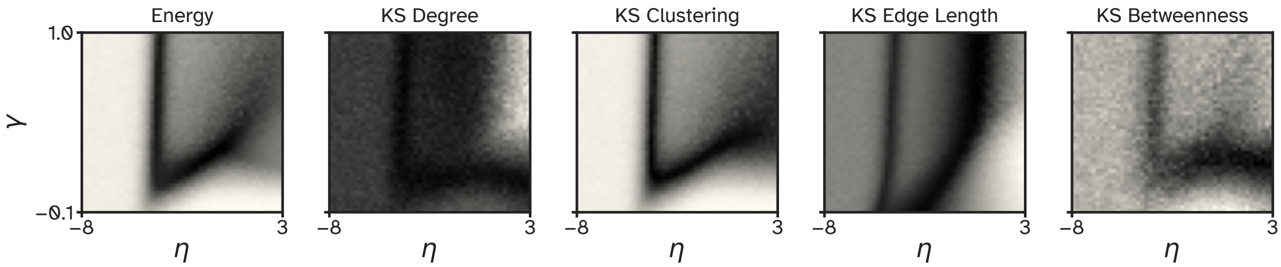


Figure S11: **Components of the KS-based energy.** Left: the landscape of the KS-based energy. Other plots: the four components that make up the KS-based energy. Lower values are darker.

The four components of the KS-based energy can be found in Figure S11. The KS-based energy is then the maximum value of the four components.

#### S4 PCA

To determine whether the distance measures share a common underlying structure in how they evaluate the networks of the morphospace, we performed principal component analysis (PCA) over the eight selected measures (see Section 4.3). This gave us the principal axes of variation shown in Figure S12. The scree plot (Figure S12A, upper plot) reveals that PC1 captures the majority of variance (55.1%), indicating substantial agreement among measures about which parameter regions produce brain-like networks. PC2 and PC3 explain 16.1% and 15.0% of the variance, respectively. The first three PC landscapes are given in Figure S12A, lower plot: Here, the 'V-shaped' landscape pattern that was already observed in Figure 3A can be observed as PC1. This suggests that despite the distinct mathematical formulations of the distance measures, the measures (except for communication correlation, the one measure with a negative PC1 loading) roughly converge on a shared notion of network similarity. PC2 (16.1% of variance) reveals an important axis of differentiation: As observable in the lower plot of Figure S12A, PC2 has sharper edges and thus adds details to the V-shape: For instance, the vertical edge at $\eta \approx -4$ is now split up into two vertical elements. The loadings of PC2 (see Figure S12C) indicate that this component splits the distance measures into two separate groups: DeltaCon and portrait divergence load in the positive direction, while the remaining six - KS-based energy and communicability correlation most strongly, then spectral (adjacency), and only weakly Frobenius, NetSimile and spectral (non-backtracking) - load in the negative direction. PC3 (see again Figure S12A lower plot) specifically provides detail in the region of high $\eta$ and $\gamma$ values - as this region is less relevant for biological networks, the loadings will not be discussed at this location.

As PC1 changes more gradually for small changes in $\eta$ and $\gamma$ than PC2, measures with relatively "high" PC1 loadings (with respect to the other measures and its PC2 loading) might capture more global structural features; and might extract exactly the shared notion of similarities that all selected measures (except for communicability correlation) show. Vice versa, measures with relatively "high" PC2 loadings (with respect to the other measures and its PC1 loading) might capture more local structural changes, and are more specific to each measure.

We summarize this trade-off per measure by the ratio $|PC2/PC1|$, i.e. the absolute loading on PC2 divided by the absolute loading on PC1. The eight ratios fall into three groups, separated by the two largest gaps in the ordering: 'Smoothers' are dominated by the shared consensus pattern: spectral (non-backtracking) (0.10), NetSimile (0.15), and Frobenius (0.19). Spectral (adjacency) (0.75) sits alone between the groups as a 'balancer', carrying both axes at comparable weight. The 'discriminators' load more strongly on PC2 than on PC1: KS-based energy (1.34), portrait divergence (1.38), DeltaCon (1.86), and communicability correlation (2.06), the last of which does not share the rough 'V-shape' landscape pattern in PC1 that the other measures show.

Together, these results show that the measures share substantial common ground - while additionally retaining complementary perspectives on network topology. The choice of picking either a smoother, balancer, or discriminator - or a combination of these - depends on the research question at hand.

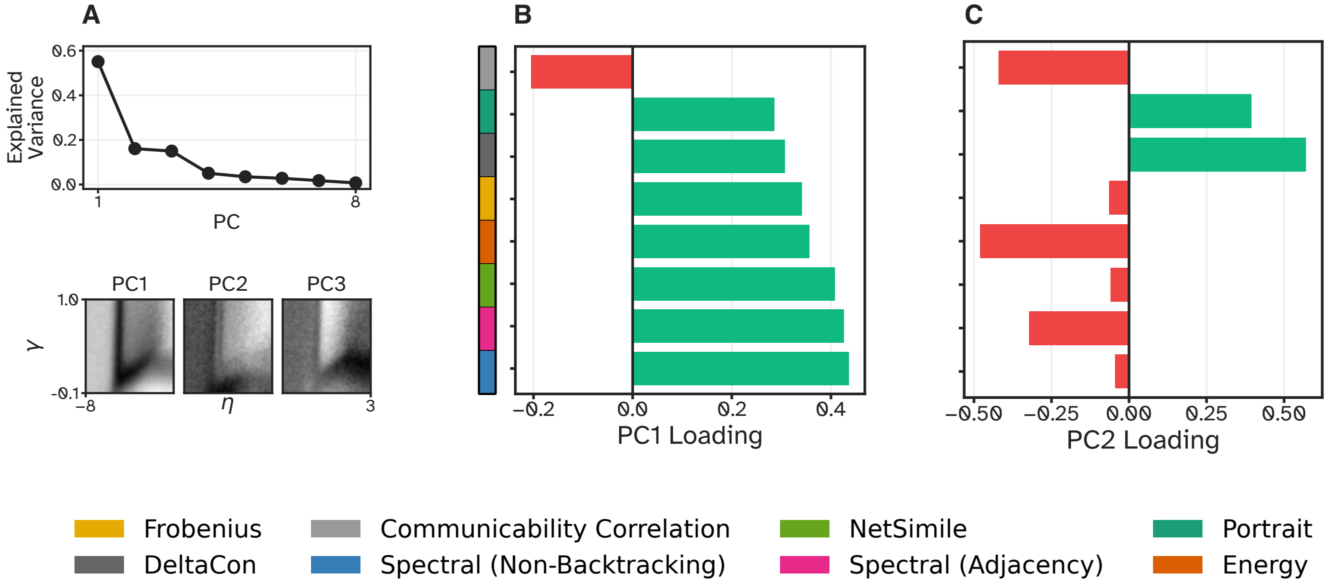


Figure S12: **Principal component analysis of the eight selected distance landscapes. A**, upper: Scree plot of the variance explained by each of the eight principal components. **A**, lower: The scores of the first three principal components mapped back onto the $(\eta, \gamma)$ morphospace, one landscape per component, with $\eta$ on the horizontal and $\gamma$ on the vertical axis. **B**: Loadings of each distance measure on PC1. **C**: Loadings of each distance measure on PC2. In **B** and **C**, the bars are in the same order in both panels, sorted by the absolute value of the PC1 loading; green bars mark positive and red bars negative loadings.

#### S5 Long-range and Interhemispheric Connectivity

To make the shape of the distance landscapes easier to read, we computed two descriptive properties of the generated networks over the same 50 × 50 grid: the long-range fraction, i.e. the share of edges longer than 90 mm - the threshold used in the main text[20] - and the interhemispheric fraction, i.e. the share of edges whose two endpoints lie in opposite hemispheres, with hemisphere membership taken from the sign of the MNI $x$ coordinate of the Schaefer-100 parcel centroids. Neither is a distance measure; both are averaged across the 10 replicates of each grid cell. Every network in the morphospace carries the same 495 edges, so both fractions are directly comparable across the grid and against the empirical consensus, which places 9.7% of its edges beyond 90 mm.

Both properties rise steeply once $\eta$ approaches and exceeds zero (Figure S13): at $\eta = 3$ the generated networks place 60.1% of their edges beyond 90 mm and 54.5% between the hemispheres, against 5.7% and 22.9% in the low-distance band at $\eta$ between −4 and −3, and both exceed what random wiring on the same nodes would give (39.9% long-range, mean edge length 80.0 mm, interhemispheric fraction 0.505). Such networks are dominated by long, cross-hemispheric connections and are thus far from the short-range-dominated empirical connectome. At strongly negative $\eta$ wiring becomes effectively random instead, so edge lengths approach the random expectation from below: once every entry of $D_{ij}^{\eta}$ is smaller than the constant (here $10^{-8}$) added to keep wiring probabilities strictly positive, adding the constant flattens the distribution and the distance term carries no information. With the offset used here this sets in gradually: 29% of node pairs sit below it at $\eta = -4$, 89% at $\eta = -5$, and 98% at $\eta = -6$; mean connection length rises from 36.8 mm at $\eta = -3.5$ to 66.2 mm at $\eta = -5.3$ and 68.9 mm at $\eta = -8$, the long-range fraction from 4.5% to 40.9%, and the interhemispheric fraction from 0.222 to 0.430

over the same range. The left arm of the V is thus the boundary at which the model stops behaving as a distance-dependent model, not a region of unusually short wiring. Only the band in between keeps wiring short and reproduces the empirical values of both properties (dashed light lines in Figure S13), and it is the same band in which the distance landscapes have their minima.

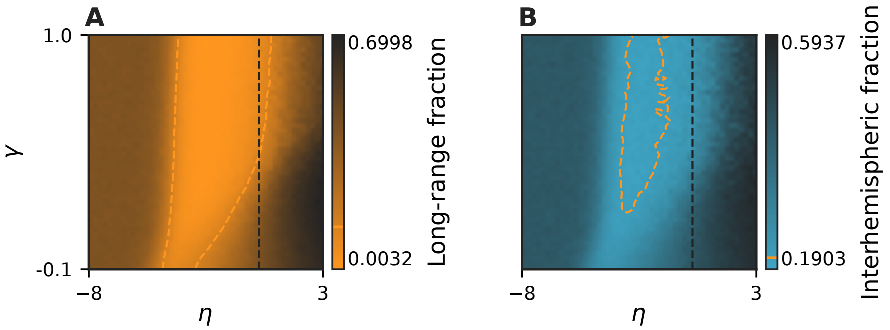


Figure S13: **Long-range and interhemispheric wiring across the morphospace.** **A**: Long-range fraction, the share of a network's edges longer than 90 mm, over the 50 × 50 grid of $(\eta, \gamma)$ combinations; each cell is averaged across the 10 networks generated at that combination. **B**: Interhemispheric fraction, the share of edges whose two endpoints lie in opposite hemispheres, on the same grid and with the same averaging; Darker cells indicate more long-range and more interhemispheric wiring, as labelled at the ends of the colour bars. The dashed orange line in each panel marks the cells whose value equals that of the empirical consensus connectome (drawn on a 3 × 3-smoothed copy of the grid). The dashed dark vertical line marks $\eta = 0$, the boundary beyond which long connections are encouraged rather than penalized.

#### S6 Hub Topography

The plausibility criterion used in the main text is based on metabolic cost (i.e. $\eta > 0$ that makes the GNM reward long connections, thus encouraging networks that are metabolically expensive and thus biologically implausible), together with a comparison of the degree and connection-distance distributions. These criteria are distributional: they constrain how many connections of a given kind a network has, but not where those connections sit. However, a network can match the degree distribution of the consensus exactly and still place every hub in the wrong region. This section asks whether the plausibility criterion can thus be sharpened into a topographic one; and finds that (on this generative model) it cannot.

A topographic criterion is only defined when the nodes of the two networks correspond to each other, which they do here: every generated network is built on the same Schaefer-100 parcellation as the empirical consensus, so node $i$ denotes the same brain region in both. Each network can then be summarised as a node-level map, assigning every node the value of one nodal quantity, so that two networks can be compared node by node rather than distribution against distribution. We use nodal degree as that quantity, giving each network a degree map.

A single realisation of a stochastic generative model is noisy, so a parameter combination is represented by the mean of the degree maps of its ten replicates. This mean map is then correlated (Pearson) with the consensus. The alternative approach - correlating each replicate separately and averaging the ten correlations - leaves the replicate noise inside every correlation and therefore scores lower: the best value generated anywhere in the $(\eta, \gamma)$-space is $r = 0.09$ per replicate against $r = 0.21$ for the alternative approach. We use the averaged maps because they are the more favourable of the two estimates, so that the negative result below cannot be attributed to replicate noise. Scoring is carried out for all 2,500 $(\eta, \gamma)$ combinations (Figure S14B).

We calibrated the criterion on the empirical data. For this, we computed the same correlation for each of the 100 individual connectomes that were used to build the consensus. The empirical degree map is (as expected) highly correlated both between (i) empirical individuals and (ii) between each individual and the consensus: (i) the mean degree maps of two disjoint halves of the subject pool correlate at $r = 0.99$, and (ii) a single individual connectome correlates with the consensus degree map at $r = 0.681 \pm 0.062$, with the weakest individual reaching $r = 0.410$. That distribution across individuals marks the noise ceiling of the criterion: In the best case, a generated and genuinely brain-like degree map could achieve this correlation value, if it would perfectly replicate a brain-like hub topography.

Between the GNM networks and the consensus network, correlation values are far lower. The best agreement attained anywhere in the parameter space is $r = 0.215$, with a grid median of −0.084. In other words: not one of the 2,500 parameter combinations is as good a predictor of empirical hub placement as the weakest individual brain (Figure S14C). Since the ceiling is out of reach everywhere in the grid, the criterion cannot separate the eight measures in any meaningful way: while differences between the criteria exist, they lie below the level at which hub topography is recovered at all. Two summaries of the grid show this: First, summarising each measure by its 100 best-fitting parameter combinations instead of only looking at the one best-fitting parameter combination, the median agreement spans $r = -0.039$ (DeltaCon) to $r = -0.119$ (Frobenius), and every measure's interquartile range lies below zero, i.e., the whole spread sits inside the noise of the landscape.

This failure of the GNMs to reproduce the empirical hub topography might be because the only node-level information that the cost term of the generative model receives is spatial centrality (i.e. the negated mean Euclidian distance from a node to all others). This may generate a hub layout that is dictated by geometry. This is reflected in the fact that across each measure's 100 best-fitting parameter combinations, nodal degree correlates with spatial centrality at a median of $r = 0.63$ (Frobenius) to $r = 0.79$ (DeltaCon), reaching $r = 0.90$ elsewhere in the grid (Figure S14D). In contrast, empirical nodal degree carries no such dependence, neither in the consensus ($r = -0.046$) nor across individuals ($r = 0.013 \pm 0.079$). Generated hubs therefore sit where spatial centrality (i.e.: geometry) puts them, whereas empirical hubs do not.

This produces the following consequence for the benchmark: As the generative model does not produce networks A distance measure can only be rewarded for finding networks that exist in the space it searches. Because no region of the homophily-based morphospace reproduces empirical hub placement, topographic agreement measures the generative model rather than the measure applied to it, and we do not use it as a plausibility axis in the main analyses. It is, however, a usable target for generative rules: the calibration above gives an explicit level to reach ($r \approx 0.68$, the individual-connectome ceiling) and an explicit confound to avoid (a degree map explained by spatial centrality), and generative rules in which nodes begin wiring at different developmental times can produce hubs that are more in topographic alignment with empirical brain networks[21].

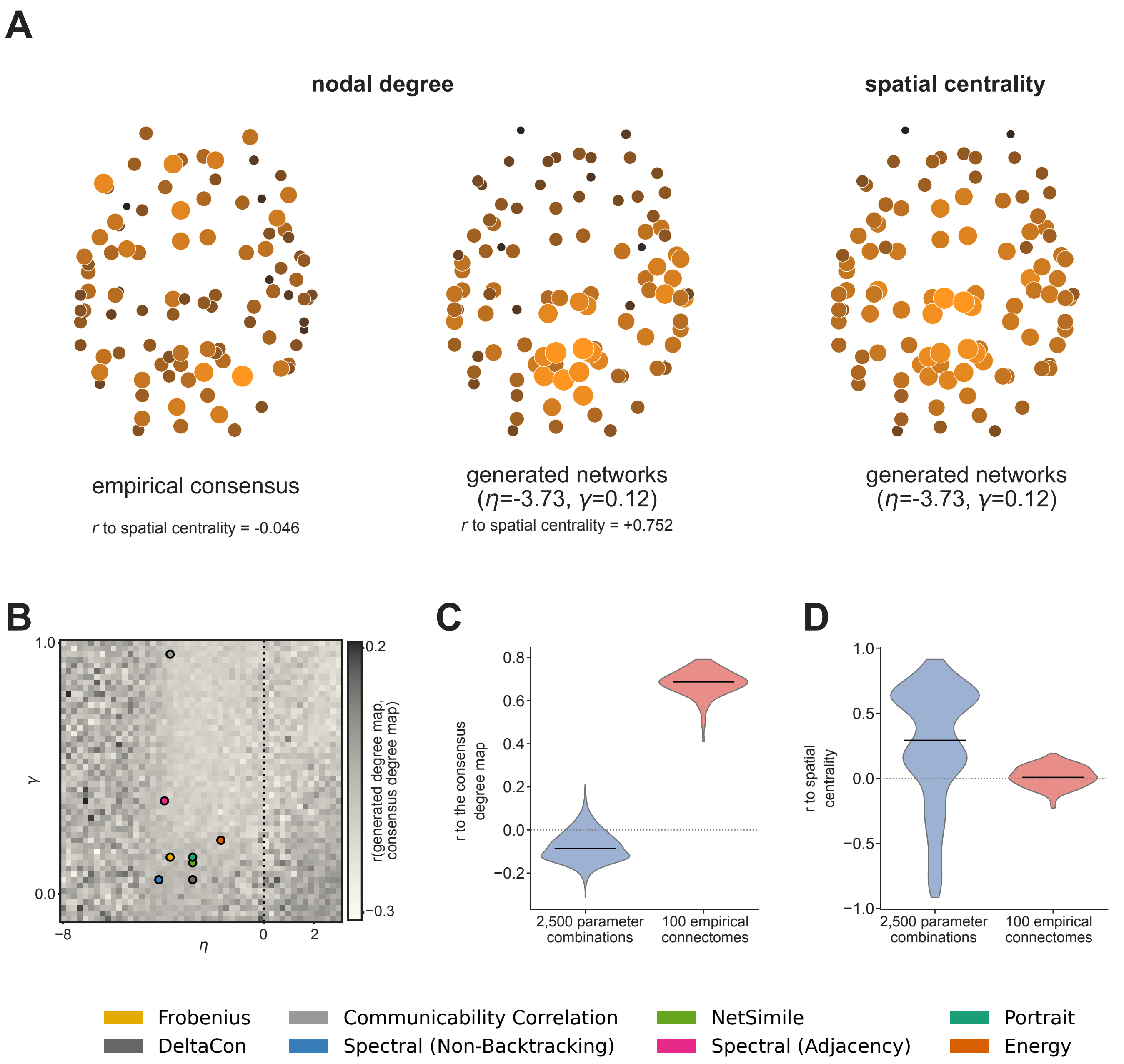


Figure S14: **Hub placement in the empirical consensus, in the generated networks, and its relation to spatial centrality.** **A**: Axial view of the 100 Schaefer nodes. Node size and colour indicate the value of the nodal degree of the empirical consensus (left), the nodal degree of the generated networks (middle), and the spatial centrality (right). The maps for the generated networks were averaged across 10 networks, that all were generated at the parameter combination defined by taking the median of the best-fitting $(\eta, \gamma)$-combinations of the eight measures, $(\eta = -3.73, \gamma = 0.12)$. Spatial centrality is defined as the negated mean Euclidean distance from a node to all others. Each map is scaled to its own range. **B**: Hub-placement agreement across the morphospace, i.e. the node-wise Pears on correlation between the repl icate-averaged degree map of each $(\eta, \gamma)$ grid cell and the degree map of the empirical consensus. Circles mark each measure's best-fitting parameter combination; the dotted line marks $\eta = 0$. **C**: Distribution of the node-wise correlation of the degree map with the consensus degree map. Blue violins span the 2,500 parameter combinations and red violins the 100 individual connectomes; black lines mark medians and the dotted line marks zero. **D**: Distribution of the node-wise correlation of the degree map with spatial centrality. The color coding is again as in **C**.

#### S7 Timing Analysis

Figure S15 shows the time needed to calculate the distance between two graphs for all measures described in this paper; precise numbers can be found in Table S2. The times were averaged across 25,000 comparisons. The figure also shows a few further measures that have been tested, but which have been dismissed early on due to unsuitability for the benchmarking task (e.g.: bad performance, high similarity to a different measure).

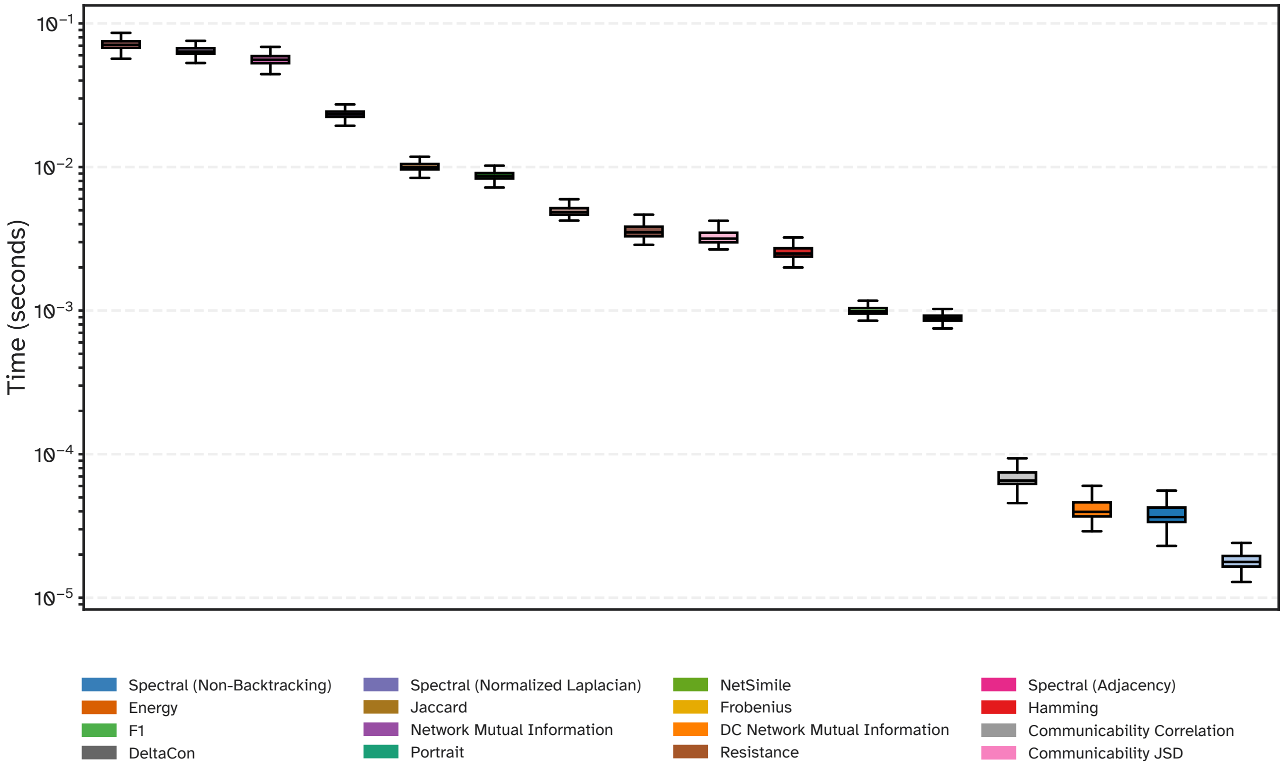


Figure S15: **Timing of all distance measures.** Computation time needed per comparison for each tested distance measure, calculated over 25,000 network comparisons (log scale).

#### S8 Synthetic Parameter Recovery

**Parameter ranges used in paper** To directly assess whether distance measures can recover the parameter combination that generated a given synthetic network (i.e.: the core task in generative network modelling research) we conducted a synthetic parameter recovery analysis. 100 parameter combinations were drawn uniformly from the whole morphospace ($\eta \in [-8, 3]$, $\gamma \in [-0.1, 1]$) as ground-truth targets, and one test network was generated at each of them. Each test network was compared against 100 grid-point consensuses on a dedicated 10 × 10 recovery grid that spans the full morphospace, where each grid point was represented by a consensus built from 30 networks. Thus, one grid step is 1.22 in $\eta$ and 0.12 in $\gamma$. The recovery error is expressed in grid steps, defined as the Euclidean distance in grid-index space between the true nearest cell and the predicted cell, and averaged across the 100 test networks. The maximum possible error is $\sqrt{9^2 + 9^2} \approx 12.73$. Figure S16 shows the true ($\times$) and recovered (•) parameter locations for all 16 measures. The ranking of the measures by mean recovery

| Method | Time (milliseconds, mean ± std) |
|---|---|
| Energy | 71.7234 ± 6.3094 |
| Spectral (Non-Backtracking) | 67.4767 ± 16.2705 |
| NetSimile | 57.2284 ± 10.6295 |
| Portrait | 23.4647 ± 2.1377 |
| DC Network Mutual Information | 10.1781 ± 0.9333 |
| Network Mutual Information | 8.7744 ± 0.8688 |
| Spectral (Adjacency) | 4.9728 ± 0.5206 |
| Spectral (Normalized Laplacian) | 3.7044 ± 3.0545 |
| DeltaCon | 3.4371 ± 3.2493 |
| Resistance | 2.6676 ± 2.1617 |
| Communicability JSD | 1.0205 ± 0.1757 |
| Communicability Correlation | 0.9046 ± 0.1741 |
| F1 | 0.0630 ± 0.0205 |
| Jaccard | 0.0478 ± 0.0567 |
| Hamming | 0.0442 ± 0.0802 |
| Frobenius | 0.0189 ± 0.0056 |

Table S2: **Timings of all distance measures.** Mean and standard deviation of the computation time per comparison, calculated over 25,000 network comparisons.

error is given in Figure 5E of the main article, and its decomposition into the two parameter axes in panel **A**. The full methodology is described in Section 4.7.

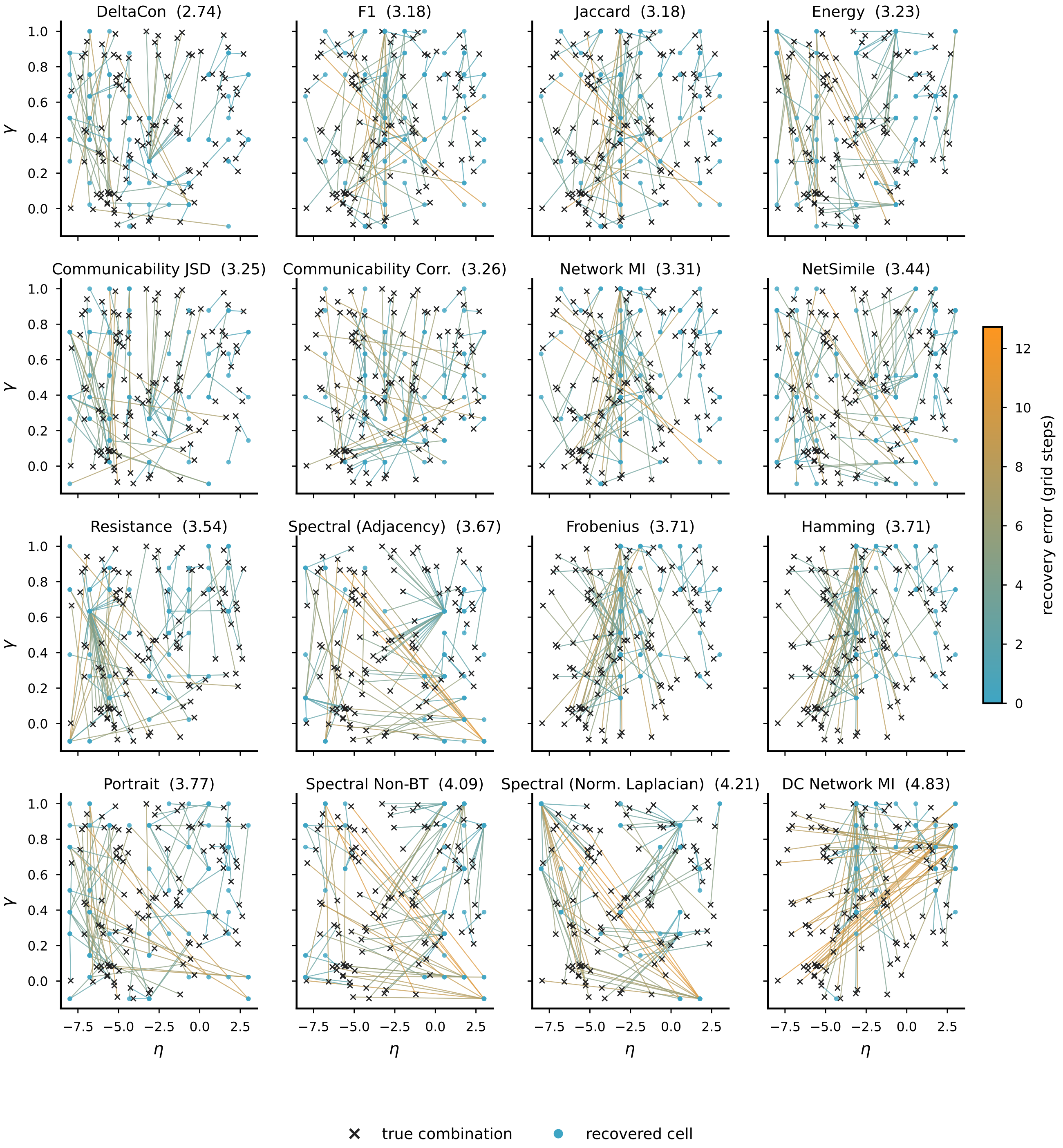


Figure S16: **Synthetic parameter recovery on the widely-spread ground-truth combinations.** Recovered locations per measure. In each of the 16 subpanels, crosses ($\times$) mark the 100 uniformly drawn true parameter combinations and filled circles (•) the cells recovered on the 10 × 10 recovery grid. Each line connects the true and predicted parameter combination for a test network, and its colour gives that network's recovery error in grid steps (colour bar). The value printed in each subpanel title is the mean Euclidean distance in grid-index space between the true and recovered cells across all 100 test networks; the maximum possible error is approximately 12.73. The subpanels are ordered by that mean.

**Parameter recovery within the plausible range for human connectomes** The recovery analysis above samples parameter combinations spanning the entire morphospace. This allows us to analyse how well a measure can discriminate coarsely between networks that are structurally very different. Empirical research, however, operates mostly inside a far narrower band: individual connectomes and species cluster near the biologically plausible combination, and the practical question is whether a measure can separate two networks generated by *similar* parameters. We therefore repeated the recovery experiment within a narrower parameter window.

We sampled 100 ground-truth parameter combinations uniformly from $\eta \in [-3.96, -1.49]$ and $\gamma \in [0.08, 0.30]$, and generated one network per combination. The window was not defined by chance, but is the pooled interquartile range along each axis of the locations of the 100 best-fitting networks of each of the eight selected measures against the empirical consensus (800 points in total); i.e., the window covers the region an empirical study would end up searching. It was re-derived on the corrected landscapes described in Section 4.2, and the ground-truth networks, the recovery grid, and every number below were regenerated inside it.

The recovery used a dedicated 10 × 10 grid spanning $\eta \in [-4.31, -1.14]$ and $\gamma \in [0.05, 0.34]$. This is slightly wider than the sampling window, so that the recovery near its edges is not truncated. Each grid point is represented by a consensus built from 20 networks.

Scoring was performed in two complementary ways: The first is identical to the wide-target experiment, where the mean recovery error is measured in grid steps, i.e. the Euclidean distance in grid-index space between the recovered cell and the cell closest to the true parameters, with a maximum of approximately 12.73 on a 10 × 10 grid. Both grids are 10 × 10, so the two experiments produce numbers on the same scale, but those numbers do not mean the same thing and we do not compare them directly: one grid step is 0.35 in $\eta$ here against 1.22 on the wide grid, so each experiment is read against its own chance level instead. Here, a measure recovering a uniformly random cell scores 4.53 grid steps (2.90 along a single axis). The second score is the Pearson correlation between true and recovered $\eta$. The two scoring functions are needed together as they complement each other's shortcomings: the grid-step error is absolute and defined for every measure, but it pools both axes, so a measure can rank well on it while tracking neither parameter closely. Correlation isolates a single axis, but it is insensitive to a constant bias or a compressed recovery range, and is undefined for measures whose recovery is constant.

Recovery inside the window is poor for every measure (Figure 5B in the main article). The best score is 2.36 grid steps (DeltaCon), followed by portrait divergence (3.37), communicability correlation (3.63), NetSimile (3.72), and communicability JSD (3.85). Five of the 16 measures score at or beyond the chance level of 4.53: resistance reaches 4.57, spectral (adjacency) 5.26, spectral (normalized Laplacian) 6.28, KS-based energy 6.29, and spectral (non-backtracking) 6.68. The two spectral measures retained in the benchmark are the ones that barely vary their answer: spectral (non-backtracking) and spectral (adjacency) each return six distinct cells for the 100 ground-truth networks, of which only four differ along $\eta$.

The per-axis decomposition shows that the difficulty is not confined to one parameter. Along $\eta$ the mean error ranges from 1.55 index steps (Frobenius, Hamming) and 1.78 (DeltaCon) to 4.76 (spectral non-backtracking) and 4.81 (spectral (normalized Laplacian)), with 11 of the 16 measures beating the per-axis chance level of 2.90. Along $\gamma$ the error ranges from 1.18 (DeltaCon) and 1.78 (NetSimile) to 4.24 (spectral (adjacency)) and 4.30 (spectral (non-backtracking)), with only six measures beating that same level. Both parameters therefore carry recoverable signal in this window, and which of the two a measure recovers better varies: Frobenius and Hamming are the most accurate on $\eta$ and among the weakest on $\gamma$, while DeltaCon is the most accurate on $\gamma$ and second on $\eta$, which is what puts it first overall. Scored on $\eta$ alone as the correlation between true and recovered values, Frobenius and Hamming lead (both 0.259), followed by F1 and Jaccard (both 0.240), communicability JSD (0.217), NMI (0.199), and communicability correlation (0.195). Lower correlations are found for DeltaCon (0.126), DC-NMI (0.103), NetSimile (0.081), and portrait divergence (0.063). KS-based energy (−0.011) and spectral (adjacency) (−0.013) are indistinguishable from zero, and resistance (−0.084), spectral (normalized Laplacian) (−0.210), and spectral (non-backtracking) (−0.279) recover $\eta$ in the wrong direction.

How far the wide-target ranking transfers into the window depends on which of the two scores is used. Scored in grid steps (Figure 5E in the main article), the correspondence is weak (Kendall $\tau = 0.19$, $p = 0.32$): most of the field changes places, with KS-based energy falling from rank 4 to 15, F1 and Jaccard from 2 to 9, and Frobenius and Hamming rising from 11 to 6, portrait divergence from 13 to 2, NetSimile from 8 to 4, and communicability correlation from 6 to 3, while spectral (non-backtracking) drops from 14 to 16. DeltaCon is the one measure that holds its position, ranking first in both regimes. Scored on $\eta$ alone, the transfer is no better ($\tau = 0.17$, $p = 0.37$, across all 16 measures, each of which has a defined correlation): DeltaCon tracks $\eta$ best on the wide targets ($r = 0.82$) but only tenth-best inside the window ($r = 0.126$), where Frobenius and Hamming lead (both $r = 0.259$) from sixth place on the wide targets ($r = 0.70$); KS-based energy holds third place on the wide targets ($r = 0.76$) and is indistinguishable from zero inside the window ($r = -0.011$). Both scores therefore tell the same story: which measure recovers parameters well depends on how far apart the candidate parameter sets are. The two window rankings agree with each other only moderately ($\tau = 0.32$, $p = 0.09$), the difference being carried by measures whose $\gamma$ estimate is accurate while their $\eta$ estimate is not, DeltaCon and portrait divergence in particular. Under either score, a measure selected on widely-spread targets is not guaranteed to be the best choice inside the range in which empirical work is done.

#### S9 Robustness of the Recovered Parameter Combinations to Reference Noise

Every fit in this work depends on the group consensus connectome as empirical reference. However, real reconstructions carry noise, as a small number of its edges could plausibly have been placed elsewhere. How much, then, does the best-fitting $(\eta, \gamma)$ combination a measure returns depend on the exact reference network it was fitted to? We rewired the consensus itself by increasing amounts, re-scored the morphospace against each perturbed reference and figured out the best-fitting $(\eta, \gamma)$ combination, and measured how far that combination moved.

This experiment differs from the rewiring analysis in the main text (Figure 4D) in both what is perturbed and what is measured. There, the consensus and its progressively rewired copy are the *pair being compared*. The read-out is the distance between them, and no generated network and no landscape is involved. The result characterises the measure's response function. In the experiment described here, the rewired consensus *replaces the reference* against which all 25,000 generated networks are scored. The read-out is the position of the minimum of the resulting landscape (i.e., their best-fitting

$(\eta, \gamma)$ parameters). The result is the change in where the best-fitting parameter combination lies. The first experiment thus asks how strongly a measure reacts to a structural change, the second asks whether the conclusion drawn from it survives a slightly different reference. The two experiments are therefore not redundant: the second depends on the geometry of the landscape (i.e. how sharp and how isolated the minimum is) and not only on the response function, so a measure can react steeply to rewiring and still hold its best-fitting combination, or react weakly and still lose it.

Starting from the consensus ($E = 495$ edges), we applied $s$ connected double-edge swaps, which preserve density, connectivity, and the exact degree sequence, and hence change only where connections sit. We used $s \in \{1, 2, 5, 10, 20, 50, 100, 200, 495\}$. As in the degeneration analysis of the main text, we report the perturbation level as the effective degeneration: the fraction of the $E = 495$ consensus edges that actually differ from the perturbed reference. This effective degeneration value is again smaller than the nominal $2s/E$, because swaps are rejected when they would disconnect the network, and later swaps can relocate an already-relocated edge. We used 10 independent perturbation draws per level for the four 'slow' measures (KS-based energy, NetSimile, spectral (non-backtracking), portrait divergence) and 20 for the 'fast' measures (all others). For each perturbed reference and each measure we re-recovered the best-fitting combination by taking the minimum of the replicate-averaged distance landscape over the same 50 × 50 morphospace used throughout, and expressed drift as the Euclidean distance in grid-index space between the perturbed and the unperturbed best-fitting combination (i.e.: the same grid-step unit as the parameter-recovery analyses). Fast measures were recovered on the full grid. Slow measures were recovered on a ±10-cell window around the unperturbed best-fitting combination; this window was widened to the full grid whenever the windowed minimum landed on the window edge at low or intermediate noise. At high noise an edge hit was recorded as a right-censored drift instead. Censoring affects the slow measures only and stayed rare: at most 11 of their $9 \times 10 = 90$ draws per measure (the fast measures have $9 \times 20 = 180$ draws and none censored). The window approach is what makes the experiment computationally affordable: a full-grid search costs 25,000 comparisons per draw against 441 × 10 = 4,410 inside the window, i.e. roughly 2.25 against 0.40 million comparisons per slow measure, which at the timings of Table S2 is about six days of single-core compute against about one day for the four slow measures together. We summarise each measure by its noise tolerance $N^*$, which we define as the largest effective degeneration at which the median drift is still within one grid step.

The measures differ by a factor of nearly 40 in $N^*$, and every one of the eight tolerates some noise (Figure S17). Communicability correlation ($N^* = 0.264$) holds its best-fitting combination exactly until a large fraction of the reference has been rewired; DeltaCon follows at 0.063, then Frobenius and NetSimile (0.029 each), spectral (non-backtracking) and spectral (adjacency) (0.015 each), and finally portrait divergence and KS-based energy (0.007 each), i.e. roughly two swapped edges of 495. At the smallest perturbation tested (a single swap moving approximately 1.5 edges) portrait divergence is the only measure whose best-fitting combination moves at all, by a median 1.1 grid steps.

The shape of the drift and the spread across draws also differ. At the levels where a measure shifts in its best-fitting parameter combination, the individual draws are almost always bimodal, with most draws holding the unperturbed best-fitting combination while a few relocate by a far distance. This is why the interquartile bands in Figure S17C reach from zero to several tens of grid steps at those levels while remaining tight elsewhere. Communicability correlation degrades discontinuously: its drift stays at exactly zero through 26% of edges moved and then jumps to a median 14 grid steps at 44%. Its unperturbed minimum sits close to the upper $\gamma$ edge ($\gamma = 0.96$), so that jump covers a large part of the $\gamma$ range. The median understates how abrupt this is: at that level the individual draws split between holding the original cell and jumping to the far side of the grid (interquartile range 0 to 19 grid steps), so the minimum does not drift towards a nearby optimum but switches between two distant ones. The remaining measures move more gradually from the first perturbations onward. At an intermediate level (14% of edges moved), the drift decomposes differently across measures: DeltaCon and NetSimile move only in $\eta$ (median 5 and 7 cells) whereas spectral (adjacency) moves almost entirely in $\gamma$ (15 cells), and portrait divergence moves in both (4.5 and 7 cells).

It is important to note that $N^*$ was more calculated for being an interesting observation, and that high $N^*$ should not be read as accuracy. For instance, Frobenius and DeltaCon are stable here partly because their landscapes are broad and, for Frobenius, nearly flat along $\gamma$, so the minimum has little reason to move. This has been shown in Section S8 already, where Frobenius tracks the true $\eta$ only weakly inside the plausible window ($r = 0.259$). Thus, a stable estimate is not automatically an informative one. The relevant insight from this experiment is the following: tolerance to reference noise depends on the measure, and both a high and low tolerance can be beneficial (for parameter combination recovery, and for the detection of edge-level changes, respectively). For instance, the field-standard KS-based energy is (together with portrait divergence) the least tolerant of small changes in the reference it is fitted to - or phrased differently, the most sensitive to edge-level changes.

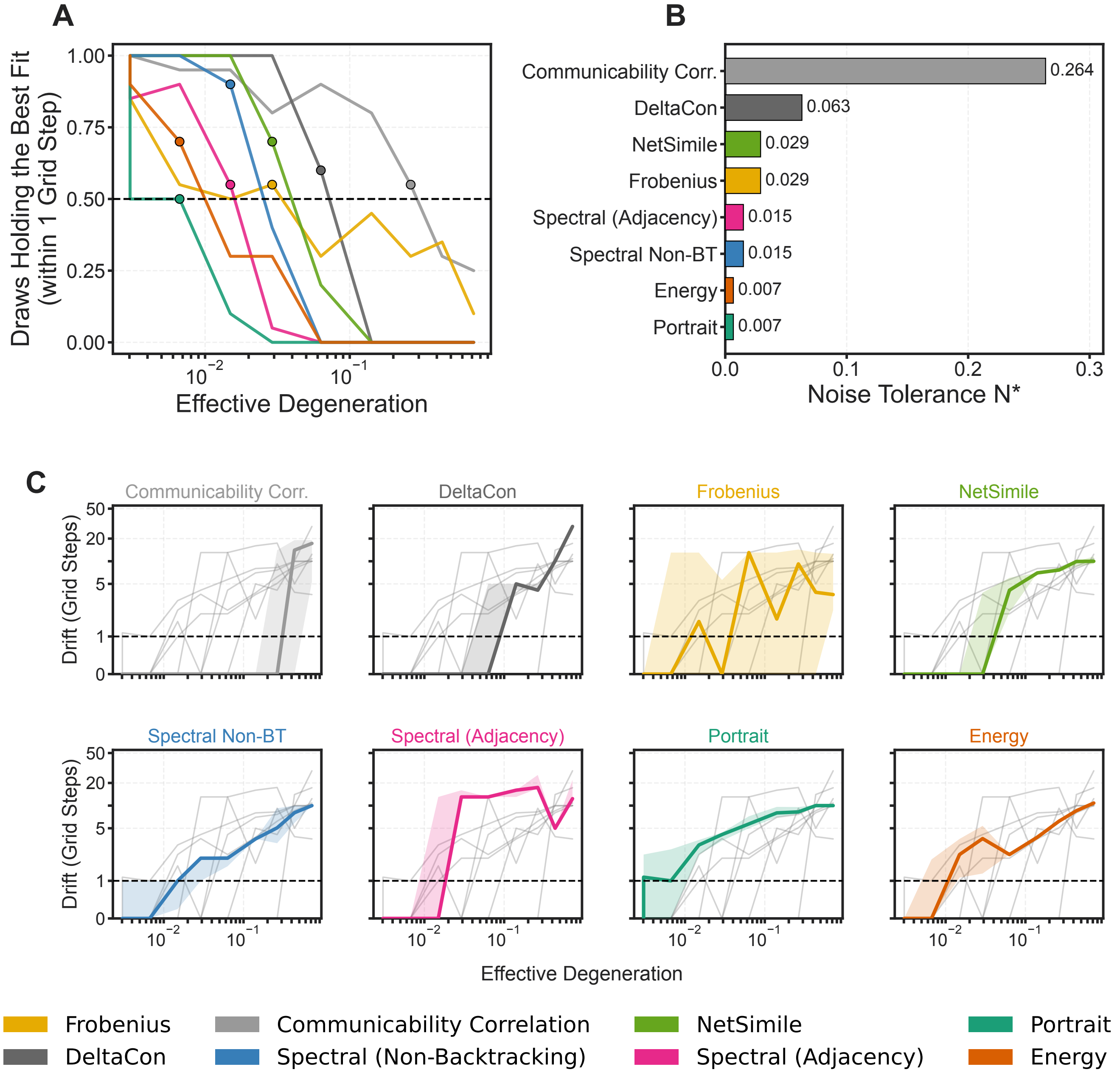


Figure S17: **Drift of the recovered parameters under progressive rewiring of the reference. A**: The fraction of perturbation draws whose recovered best-fitting $(\eta, \gamma)$ combination is still within one grid step of its unperturbed location against the effective degeneration. Effective degeneration is again defined as the fraction of the $E = 495$ consensus edges that differ from the perturbed reference, averaged over the draws at each swap count (log scale). The dashed horizontal line marks half the draws; the filled circle on each curve marks that measure's noise tolerance $N^*$. **B**: Noise tolerance $N^*$ per measure, i.e. the largest effective degeneration at which the median drift remains within one grid step, with the value printed beside each bar. **C**: Median drift from the unperturbed location in grid steps. The shaded band spans the interquartile range across the perturbation draws at each level. The medians of the other seven measures are shown in grey. The vertical axis is linear below one grid step and logarithmic above it, so that the region in which $N^*$ is read off stays resolved. The dashed horizontal line marks the one-grid-step threshold. Panels are ordered by $N^*$.

[9] Michele Berlingerio, Danai Koutra, Tina Eliassi-Rad, and Christos Faloutsos. NetSimile: A Scalable Approach to Size-Independent Network Similarity, September 2012.

[10] Danai Koutra, Neil Shah, Joshua T. Vogelstein, Brian Gallagher, and Christos Faloutsos. DeltaCon: Principled Massive-Graph Similarity Function with Attribution. *ACM Trans. Knowl. Discov. Data*, 10(3):28:1–28:43, February 2016.

[11] Dan Chen, Dan-Dan Shi, Mi Qin, Si-Meng Xu, and Gui-Jun Pan. Complex network comparison based on communicability sequence entropy. *Physical Review E*, 98(1):012319, July 2018.

[12] Nathan D. Monnig and Francois G. Meyer. The Resistance Perturbation Distance: A Metric for the Analysis of Dynamic Networks, August 2017.

[13] helcio. Hfelippe/network-MI, July 2025.

[19] Zhen-Qi Liu, Vincent Bazinet, Justine Y. Hansen, Filip Milisav, Andrea I. Luppi, Eric G. Ceballos, Asa Farahani, Laura E. Suarez, Golia Shafiei, Ross D. Markello, and Bratislav Misic. Netneurotools: A trainee-oriented approach to network neuroscience, September 2025.

[20] Stuart Oldham, Alex Fornito, and Gareth Ball. Coming up short: Generative network models fail to accurately capture long-range connectivity, November 2024.

[21] Francesco Poli, Stuart Oldham, Alexia Mousley, Edward T. Bullmore, Petra E. Vértes, and Duncan E. Astle. Right time, right place: Heterochronicity shapes brain network formation, October 2025.
